# Can Protein Structure Prediction Methods Capture Alternative Conformations of Membrane Proteins?

**DOI:** 10.1101/2023.08.04.552045

**Authors:** Tengyu Xie, Jing Huang

## Abstract

Understanding the conformational dynamics of proteins, such as the inward-facing (IF) and outward-facing (OF) transition observed in transporters, is vital for elucidating their functional mechanisms. Despite significant advances in protein structure prediction (PSP) over the past three decades, most efforts have been focused on single-state prediction, leaving multi-state or alternative conformation prediction (ACP) relatively unexplored. This discrepancy has led to the development of highly accurate PSP methods such as AlphaFold, yet their capabilities for ACP remain limited. To investigate the performance of current PSP methods in ACP, we curated a dataset, named IOMemP, consisting of 32 experimentally determined high-resolution IF and OF structures of 16 membrane proteins. We benchmarked 12 representative PSP methods, along with two recent multi-state methods based on AlphaFold, against this dataset. Our findings reveal an escalating bias towards one specific state in deep learning-based methods and a remarkably consistent preference for specific states across various PSP methods. We elucidated how coevolution information in MSAs influences the state preference. Moreover, we showed that AlphaFold, when excluding coevolution information, estimated similar energies between the experimental IF and OF conformations, indicating that the energy model learned by AlphaFold is not biased towards any particular state. Our IOMemP dataset and benchmark results are anticipated to advance the development of robust ACP methods.

## 1 Introduction

Proteins, as dynamical entities, continuously evolve their conformations in response to their biophysically functional states and physiological environments. This is particularly evident in transmembrane proteins, such as transporters, which can adopt well-defined inward-facing (IF) or outward-facing (OF) conformational states. Robust exploration and accurate determination of protein conformational changes provide essential knowledge for gaining mechanistic insights into their functional modes [1]. Targeting alternative conformations is also a potential strategy for achieving high efficacy and selectivity for drug design. However, the conformational ensemble populated by a protein exhibits a highly complex landscape, and employing experimental techniques or computational methods for high-throughput determination of key alternative conformations remains challenging. The recent success of protein structure prediction (PSP) methods for single-state structures have been revolutionizing the broad field of biological science [2, 3]. There is thus both an urgent need and significant promise in substantially improving alternative conformation prediction (ACP).

In the past 30 years, the development of multiple-state PSPs has lagged behind that of single-state PSPs. A key reason for such disparity is the extremely limited availability of multiple conformational states resolved by structural biology experiments, compared to the abundance of experimentally-resolved protein structures for single-state PSPs. For example, only 96 structurally validated fold-switching proteins, whose alternative conformations involve the transition of secondary structures, have been identified from the Protein Data Bank (PDB) [4]. For transporters, a previous study reported the IF and OF conformations of merely 59 proteins obtained through homologous modeling, with structural qualities not yet fully scrutinized [5]. Consequently, the scarcity of experimental data on multiple conformational states hinders the training and generalization of data-driven methods, as well as the benchmark of ACP methods.

Despite the limited multiple-state data from experiments, researchers have proposed several multiple-state structure prediction methods using AlphaFold as a basic tool [6, 7, 8, 9, 10]. Del Alamo et al. proposed to reduce the depths of input multiple sequence alignments (MSAs) through stochastic subsampling to drive AlphaFold to sample alternative conformations (termed AF-depth hereafter) [6]. Wayment-Steele et al. introduced AF-cluster [7], hypothesizing that coevolutionary signals of multi-states can be deconvolved by clustering sequences. Furthermore, the SPEACH AF method [8] samples alternative conformations through alanine mutagenesis to bias the structure models generated by AlphaFold. Apart from AlphaFold-based multiple-state methods, Janson et al. designed idpGAN, a generative adversarial network trained on the coarse-grained representation of molecular dynamics (MD) trajectories of intrinsically disordered proteins, for predicting their conformational ensembles [11]. Due to the absence of large enough or standardized datasets of alternative conformations as gold standards, these multiple-state methods were validated on small and disparate protein test sets, which complicates the direct evaluation of these methods and comparisons of their performance. There is thus an urgent need to expand and standardize the datasets for alternative conformations to advance the field.

As for PSP, MSA lies at the center of most ACP methods. Techniques such as AF-depth, AF-cluster, or SPEACH AF can be considered as different ways to subsampling or masked utilization of MSAs. They all pivot around the central hypothesis that the co-evolution information embedded in MSAs includes the structural information of alternative states, which could potentially be retrieved through manipulating the MSA inputs in AlphaFold. The hypothesis is supported by a recent study demonstrating that protein fold-switching can be achieved through amino acid substitutions in an evolutionary pathway, as evidenced by observations in homologous sequences [12]. Coevolution has been investigated for three decades in the context of PSP, analyzed at scales ranging from coarse to refined, including contact maps, residue pair distances, and 3D structures [13]. Despite some early emphasis on the presence of alternative conformations in coevolutionary information [14, 15], most coevolution-based PSP methods predominantly focus on single-state structure prediction. Given the emerging need to adopt PSP methods for alternative conformation prediction, a systematic review and testing of PSP methods from an ACP perspective can provide insights into how coevolution information is processed.

In general, PSP methods can be classified into four categories according to their strategies for utilizing MSAs, which will be introduced here in chronological order. The earliest methods applied correlation analysis directly between residue pairs, including the McLachlan Based Substitution correlation (McBASC) [16], the Mutual Information (MI), and the Mutual Information combined with averaged product correction (MIp) [17] methods. The second category comprises statistical-model-based methods that employ statistical models to identify direct coupling residue pairs from MSAs with the aim of mitigating the phylogenetic effect. Examples include the mean-field approximation direct-coupling analysis (mfDCA) [18], the pseudolikelihood maximization procedure (plmDCA) [19], GREMLIN [20], CCMpred [21], and PSICOV [22].

The introduction of deep learning (DL) marked a significant step in PSP development, forming the third category. These methods trained deep neural networks (DNNs) to infer contact maps or distance distributions from MSAs. RaptorX-contact [23] [24], ResPRE [25], trRosetta [26], and RaptorX-3DModeling [27] utilize residual neural networks (ResNNs) to effectively extract distant genomic restraints. In addition to predicting contacts or distance distributions, trRosetta and RaptorX-3DModeling further enhance their predictive capabilities by estimating interresidue orientations (angles and dihedrals). Meanwhile, the early version of AlphaFold [2] and RoseTTAFold [28] use the attention-based Transformer architectures to capture the inherent sequence-structure relationships of proteins. We note that the current version of AlphaFold (also called AlphaFold2) [3], infers protein structures directly from MSAs without using contact maps as intermediate information, although the contact maps can still be output in the form of distance distribution when required.

The last category of methods is implicitly dependent on MSAs as they use a single target sequence as input. These methods derive residue-level embedding and residue-residue couplings from unsupervised large language models (LLMs), as exemplified by ESM-1b [29], ESMFold [30] and OmegaFold [31]. We note that ESM-1b refers to the protein LLM, while ESMFold was obtained through further supervised learning using experimental protein structures. For the final two categories of DL-based methods, it’s particularly interesting to evaluate whether training with native structures deprives the capacity of MSA processing to extract information on alternative conformations.

Can these PSP methods capture alternative conformations of proteins well? In this work, we limited our investigations on membrane proteins. Membrane proteins undergo functionally-relevant conformational changes that are challenging to capture, which involve only rearrangement of TMs. Furthermore, these conformational changes are biologically important for membrane proteins to function so they should be strongly represented in coevolution information. Unlike many proteins that populate an ensemble of conformations, membrane proteins typically adopt one of the two dominant conformations, thus largely simplifying the analysis.

To this end, we first curated a dataset of the IF and OF states of membrane proteins (IOMemP), comprising 16 high-quality pairs of structures from the PDB database. We then benchmarked 12 PSP methods on the IOMemP dataset, with standardized input MSAs using the AlphaFold protocol to ensure the consistency. The predicted contact maps from these 12 PSP methods were then analyzed by comparing to the ground truth of alternative states in IOMemP. Our results interestingly reveal a consistent preference for one conformational state. We also tested two methods that generate conformational ensembles, AF-depth and AF-cluster, on IOMemP by directly comparing their predicted structures to the alternative conformations in the dataset. Inspired by a recent theoretical work suggesting that AlphaFold has learned an accurate energy function and that the MSA input of AlphaFold is used to help solve the global search problem [32], we computed the proposed energy scores of alternative states, and observed similar energies between IF and OF states. Finally, we examined AlphaFold’s preference for a state when its input is configured with solely MSAs, or IF and OF structure as templates or their combination. In summary, we present in this manuscript a curated dataset for benchmarking ACP results and explore the ability of various PSP methods in tackling the ACP problem for membrane proteins, which could enhance the understanding and facilitate the advancement of protein structure prediction methodologies.

## 2 Methods

### 2.1 Benchmark dataset construction

To evaluate the performance of current PSP methods for ACP, a dataset was curated from the PDB. It consists of 32 experimental structures representing 16 membrane proteins, each with both an IF and an OF structure. The construction of this dataset invovles the following steps.

#### 2.1.1 Filtration of membrane proteins

Our hypothesis is that experimental structures with over 70% sequence identity belong to the same protein and might represent different conformational states of that protein. As of October 13, 2021, clustering at 70% sequence identity in the PDB by MMseqs2 resulted in 56,823 proteins [33]. To recognize membrane proteins, we utilized the sequence-based topology prediction method, TOPCONS [34], yielding 2,719 candidate proteins. Out of these, 2,124 proteins had more than one structure available. To further mitigate the risk of false positives introduced by sequence-based topology prediction, we cross-referenced with a structure-based membrane orientation prediction tool [35] and required the total number of residues between the two predicted membranes (where z-coordinates of C*α* atoms range from -15 Å to 15 Å) to be larger than 60. This ensured the existence of multiple long TM regions, and reduced the set to 1,098 membrane proteins.

#### 2.1.2 Filtration of alternative states

Next, we identified membrane proteins with distinct IF and OF states, as reflected in significant structural differences in the TM regions. To compute these differences, we aligned two structures for paired residues by TM-align [36], which is capable of protein structure alignments with varying sequences. We excluded proteins with less than 80 aligned residues in TM regions, resulting in a list of 860 proteins. We note that we only considered the aligned residues in TM regions in our benchmark. We then defined an intra-distance difference score Δ*D* between two structures as follows:

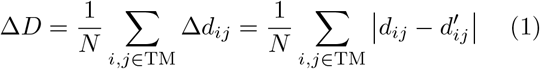

where *i, j* are indices of two residues in TM regions, *d*_*ij*_ and *d′* _*ij*_ are the distances between the *Cβ* atoms (*Cα* for glycine) of the *i*th and the *j*th residues for a structure and its counterpart, respectively. *N* is the total number of residue pairs.

For each of the 860 proteins, we calculated Δ*D* between all pairs of structures to measure the conformational differences in the TM regions, and recorded the largest Δ*D* (Δ*D*_max_) and its corresponding pair of structures. Ranking the 860 proteins by Δ*D*_max_ in descending order revealed that only a small number (63) of proteins had a Δ*D*_max_ larger than 2.0 Å, suggesting that experimental structures capturing alternative conformations are scarce (Fig. S1). Following a manual inspection and excluding structures with resolutions larger than 3.5 Å, we selected 16 top-ranked proteins with two distinct structures to form the IOMemP dataset. Detailed information on the IF and OF structures are listed in Table 1 and Data S1. The 16 proteins are transporters with a minimum of 6 TM helices. Notably, 12 out of the 16 proteins exhibit substrate efflux functionality. For proteins located on the mitochondrial membrane such as AAC3, we refer to the matrix-open conformation as the IF state and the cytoplasmic-open conformation as the OF state. We note that this study considers only the single chain of a protein.

**Table 1:**
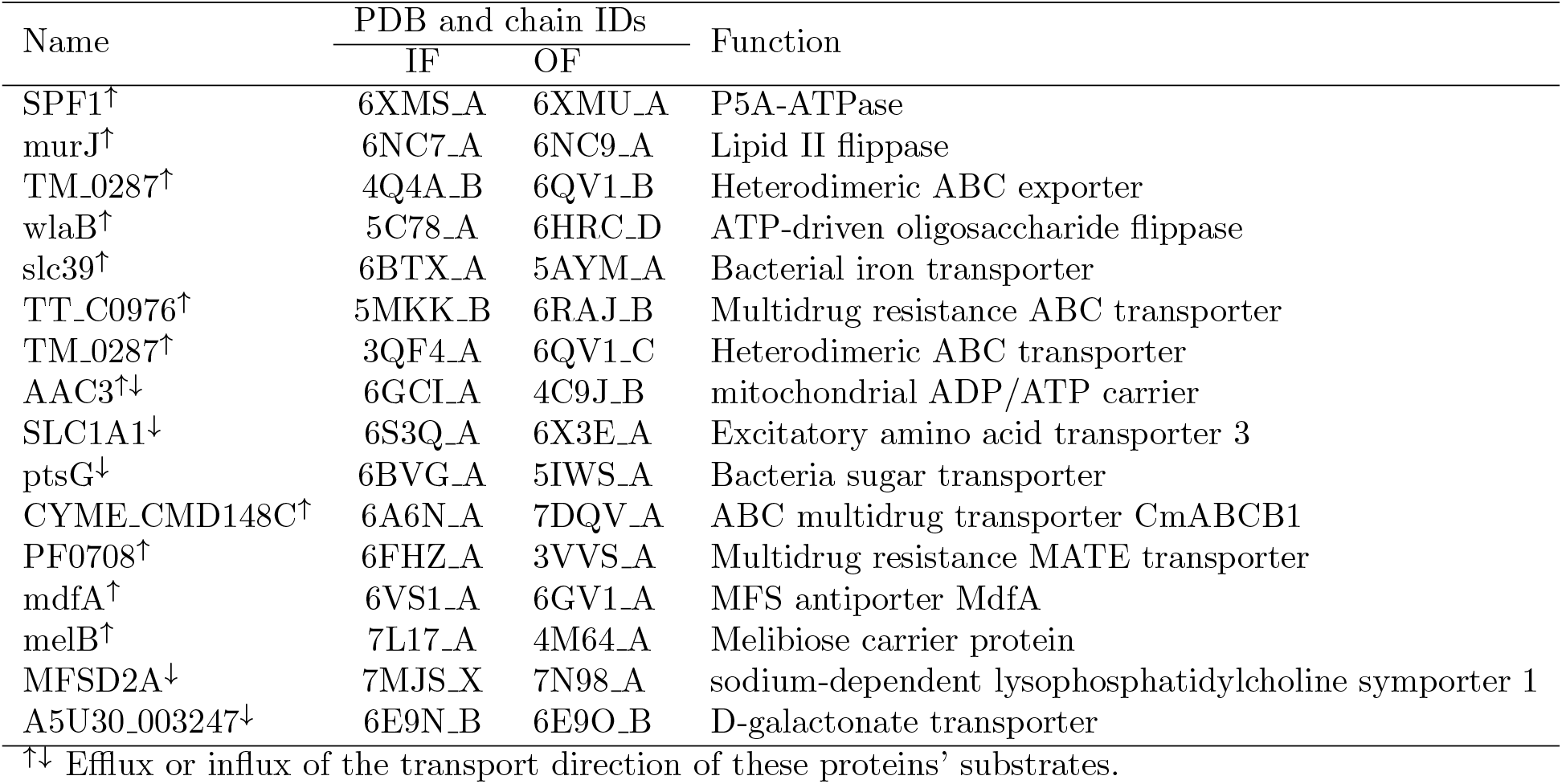
The IF and OF structures of the 16 proteins in IOMemP.

#### 2.1.3 Experimental contact maps

We defined two residues as forming a contact when the distance between their *Cβ* atoms (*Cα* for glycine) is within 8 Å in a structure. Consequently, for the 16 proteins in IOMemP, each with an IF and OF conformation, we derived 16 pairs of contact maps. These contact maps represent the ‘ground truth’ for the ACP benchmark. Based on the occurrence of contacts in both the IF and OF states, we defined the native contacts 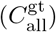 as those occurring in any of the two conformational states. These can be further divided into three subsets, contacts shared in the two states as shared contacts 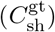 and those only in one of the states as state-specific contacts 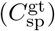, including IF contacts 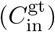 and OF contacts 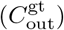. Contacts are classified into three categories according to the sequence distance between two residues: long-range (⩾24), medium-range ([12, 23]), and short-range ([6, 11]). All-range contacts (⩾6) are also considered, which encompass all the three categories. Table S1 lists the average number of different contacts with respect to the length of the sequence (*L*). Long-range contacts predominate, constituting 94% of all-range contacts, while medium-range and short-range contacts represent 4% and 2%, respectively. In the long-range contacts, shared and state-specific contacts account for 72.2% and 27.8%, respectively. Analysis will primarily focus on long-range contacts, as they effectively characterize the global structures.

### 2.2 Benchmarked PSP methods

In this work, we benchmarked 12 PSP methods: MI, mfDCA, PSICOV, CCMpred, plmDCA, RaptorX-Contact, ResPRE, trRosetta, RaptorX-3DModeling, ESM-1b, RoseTTAFold, and AlphaFold. Their corresponding default settings were used and Detailed information on these methods can be found in Data S2. Each method’s input MSAs were generated using the AlphaFold protocol, detailed in Section 2.3. These MSAs are derived from extensive sequence databases, representing the most comprehensive coevolution information available. For ESM-1b, we used the pretrained model “esm1b t33 650M UR50S” for the benchmark, which was developed without incorporating experimental protein structures [29]. For end-to-end PSP methods such as AlphaFold and RoseTTAFold, they were made to output intermediate distance distributions which can then be converted into contact maps using an 8 Å cutoff. The predicted contact maps were compared to the contact maps extracted from the experimentally determined alternative structures. After evaluating the 12 PSP methods using the most comprehensive coevolution information, we further expanded out benchmark to include two multi-state techniques, AF-depth and AF-cluster. These methods modify the initial MSAs through subsampling and clustering, respectively, resulting multiple MSA subsets. Each subset is subsequently subjected to AlphaFold prediction, yielding multiple structures that potentially represent the protein conformational ensembles.

### 2.3 Coevolution generation

To ensure the comparison between the 12 PSP methods, we uniformly generated the input coevolution following the AlphaFold protocol [3], avoiding any bias that might arise from using different MSA methods. We employed jackhammer [37] to search the UniRef90 [38] and the MGnify [39] databases, and HHblists [40] to search the BFD database. We note that this deviates slightly from the default AlphaFold practice where HHblists searchs BFD+Uniclust30 [41]. This change was necessary to avoid system crashes encountered when searching BFD and Uniclust30 simultaneously for some sequences in this study. The MSAs obtained from both jackhmmer and HHblits searches were then combined and formatted to meet the specific input requirements of each benchmarked method (e.g., converting ‘a3m’ to ‘aln’). Considering the sequence differences between the IF and OF structures, we generated two MSAs for the two states of a protein, which have nearly identical numbers of effective sequences (NEFF) as illustrated in Fig. S2.

### 2.4 Evaluation metrics

#### 2.4.1 Comparison of predicted contact maps to the IF and OF contacts in IOMemP

From a predicted contact map that consists of the contact probabilities of all residue pairs, typically the top L/*n* (*n*=1, 2, or 5) contacts are considered as the predicted contacts, after ranking the contact probabilities in descending order. Here, L denotes the number of paired residues in the TM region. When compared to the native contacts 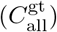, the precision of the contact map is defined as:

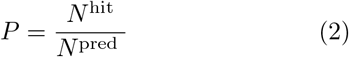

where *N* ^pred^ is the number of predicted contacts (*N* ^pred^ = L*/n*) and *N* ^hit^ is the number of predicted contacts matching the native contacts 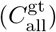.

To assess the performance of the 12 PSP methods in predicting shared and state-specific contacts, we calculated the corresponding percentage (*p*_sh_ and *p*_sp_) and coverage (*c*_sh_ and *c*_sp_), which are defined as follows:

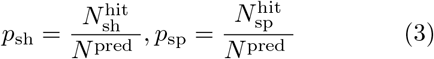

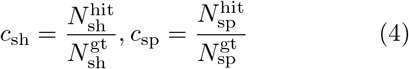

where 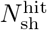 and 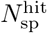 are the number of predicted contacts matching the shared contacts 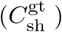 and state-specific contacts 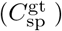, respectively. 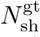 and 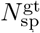 represent the numbers of contacts in 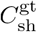 and 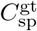, respectively. Similarly, to assess the performance in predicting the IF and OF contacts, the corresponding percentage (*p*_in_ and *p*_out_) and coverage (*c*_in_ and *c*_out_) are defined as follows:

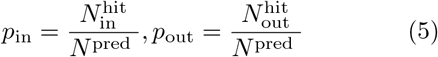

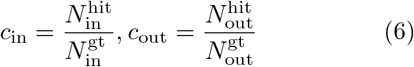

where 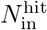 and 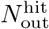 are the number of predicted contacts hitting the IF contacts 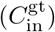 and OF contacts 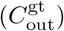, respectively. 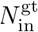 and 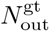 are the numbers of contacts in 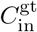 and 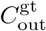, respectively.

The higher the percentage or coverage, the better the method performs in predicting the corresponding contacts. The sum of IF and OF percentages (*p*_in_ and *p*_out_) equals the state-specific percentage (*p*_sp_), while the sum of shared and state-specific percentages (*p*_sh_ and *p*_sp_) is equal to the precision (*P*) of the contact map. In this work we primarily present results using the top L contacts as similar results were obtained with top L/2 or top L/5 contacts. It’s also worth noting that for one protein, a PSP method will predict two contact maps. Both maps will be compared to the experimental contact maps, resulting in two sets of percentage and coverage.

#### 2.4.2 Preference over one state

When the two predicted contact maps generated by a PSP method both display a higher percentage to a specific state (either IF or OF), we designate this state as the preferable one for the method. The other state is considered as the unpreferable state. If this condition doest not hold, we conclude that the method does not show preference over either state.

The degree of preference (*d*_percentage_ or *d*_coverage_ of a PSP method is defined as:

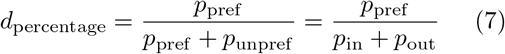

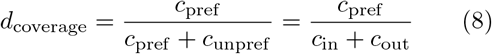

where *p*_pref_ and *p*_unpref_ represent the percentage of a predicted contact map with respective to the preferable and the unpreferable states, respectively. *c*_pref_ and *c*_unpref_ represent the coverage for either state. A higher degree of preference (*d*_percentage_ or *d*_coverage_) indicates a stronger tendency for a method to prefer a specific conformational state.

### 2.5 Estimating the energies of alternative states using AlphaFold

In addition to using coevolution as the input to test the state preferences of DL-based PSP methods, we were also interested in their state perference in the absence of coevolution information. Inspired by a recent work of Roney and Ovchinnikov [32], we computed the confidence score, which can be considered as an estimator for the energy of a particular conformational structure of a given protein sequence. In our case, we have two experimental structures in two states for one protein. One of the structure-corresponding sequences was used as the target sequence for AlphaFold, while one of the experimental structures was used as the template for AlphaFold. Upon outputting a structure, AlphaFold provides the corresponding “predicted TM-score” (pTM) and “predicted Local Distance Difference Test” (pLDDT) score [42], based on which the “composite confidence score” (*s*) can be computed as [32] :

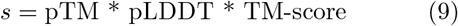

where the TM-score is calculated using TM-align and represents the structural similarity between the AlphaFold output structure and the input template [36]. A higher *s* score suggests the template structure is more favorable in energy, or more accurately in the free energy of folding.

For a specific target sequence that corresponds to either the IF or the OF structure, energies of the two alternative structures (each used separately as the template of AlphaFold) were estimated. A large difference in the energy score suggest strong preference for a particular state. We note that before assigning the coordinates of the template to the features of the target, we need to align the seqeunces of the target and the template. This step, which is not taken into account in the previous work [32], ensures meaningful calculation of *s* when the template sequence differs from the target sequence.

## 3 Results

### 3.1 PSP methods are performing increasingly well in predicting shared contacts

Fig. 1 compares the percentage and coverage of the top L long-range contacts, averaged over all structures in the IOMemP dataset, for the 12 PSP methods we benchmarked. These methods are arranged chronologically, with the first five methods being non-DL-based and the latter seven being DL-based. ESM-1b stands out as an unsupervised learning approach, while the remaining DL-based models were trained using native protein structures. Among this group of supervised DL-based methods, RaptorX-Contact, ResPRE, trRosetta, and RaptorX-3DModeling employ the ResNN architecture while RoseTTAFold and AlphaFold utilize the Transformer architecture.

**Figure 1:**
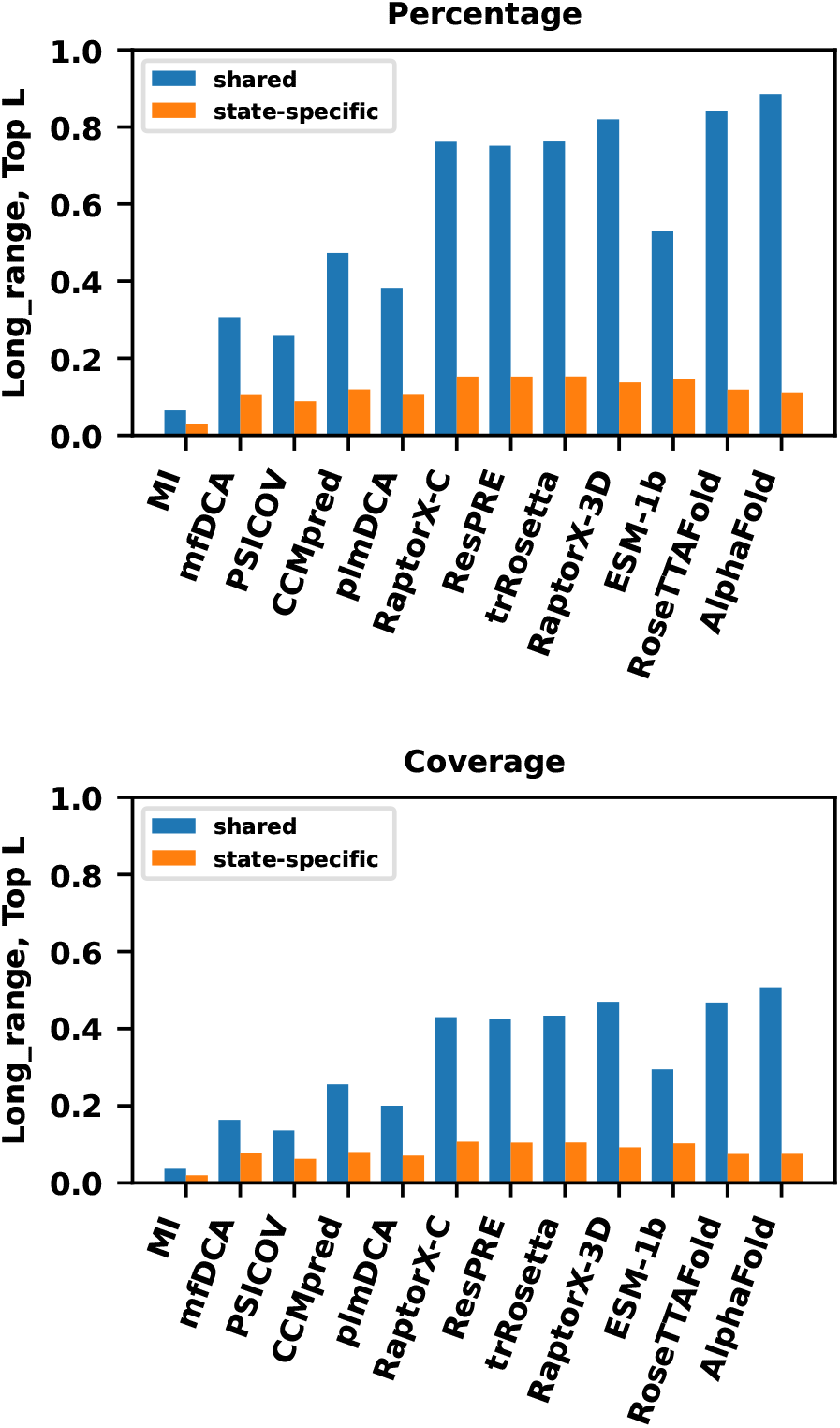
The averaged percentages or coverages of the 12 PSP methods for the top L long-range contacts. The blue bars are metrics for shared contacts and the orange bars are those for state-specific contacts (averaged on *p*_in_ and *p*_out_ or *c*_in_ and *c*_out_). “RaptorX-C” and “RaptorX-3D” refer to “RaptorX-Contact” and “RaptorX-3DModeling”, respectively.

As illustrated in Fig. 1, for shared contacts, MI has the lowest percentage and coverage while AlphaFold achieves the highest. Among the non-DL-based methods, CCMpred has the highest percentage and coverage for both shared (0.473 and 0.256) and state-specific (0.119 and 0.080) contacts. In the case of ResNN-based methods, their percentages or coverages surpass those of non-DL-based methods, in particular for shared contacts. Of the four ResNN-based methods, RaptorX-3DModeling slightly outperforms the others, with percentage and coverage reaching 0.820 and 0.470 for shared contacts. Transformer-based methods further outperform ResNN-based methods in terms of shared contacts, with AlphaFold’s percentage and coverage reaching 0.886 and 0.507. While these results were obtained considering the top L long-range contacts, the same conclusion can be drawn when using the top L/2 or top L/5 long-range contacts (Fig. S3). In addition, supervised DL-based methods also achieve high coverage for medium-range and short-range contacts (Fig. S4 and S5). The performance of all 12 PSP methods for all-range contacts is similar to that for long-range contacts (Fig. S6), as long-range contacts constitute the majority of all-range contacts (Table S1).

In terms of the average precision (*P*) of the predicted contact map, defined in this work as the sum of shared and state-specific percentages, extremely high values were obtained for most recent PSP methods, such as RoseTTAFold (*P* = 0.961) and AlphaFold (*P* = 0.998), as shown in Fig. S7. We note that our definition of *P* deviates from the common one for a single structure since we consider both IF-specific and OF-specific contacts. If we use only the contacts derived from either the IF or the OF structure, the precision drops to 0.923 and 0.953 (*p*_sh_ + *p*_in_) and 0.881 and 0.931 (*p*_sh_ + *p*_out_) for RoseTTAFold and AlphaFold, respectively (Table S2). This difference underscores the importance of considering alternative conformations in protein structure prediction. We also note that as precision increases for these methods, the percentage of shared contacts also increases in a linear relationship with precision (Fig. S8). Thus, the performance of a PSP method traditionally gauged using precision can be mirrored by the percentage of shared contacts in ACP.

### 3.2 DL-based PSP methods prioritize shared contacts over state-specific ones

As can be observed in Fig. 1 and S3, the percentage and coverage of state-specific contacts have not witness significant improvements with more recent methods, in contrast to shared contact. This becomes more evident in Fig. 2, where the correlation between shared and state-specific percentages and coverages is plotted.

**Figure 2:**
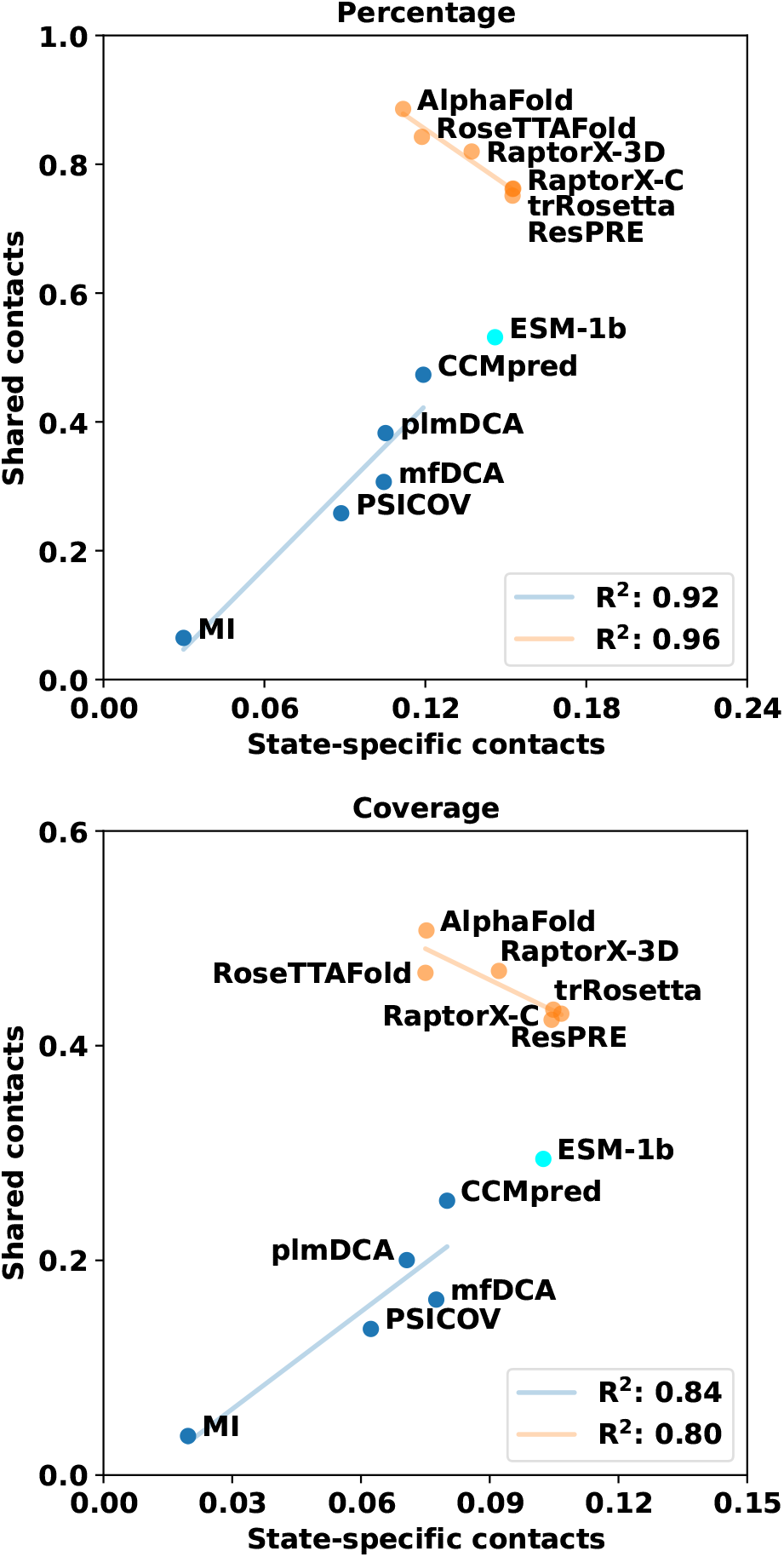
The shared and state-specific percentages or coverages for top L long-range contacts for the 12 PSP methods. The non-DL-based methods are colored in blue and the supervised DL-based methods in orange and ESM-1b in cyan. The R^2^s are the goodness of the linear regression (lines in light colors) for the non-DL-based methods and the supervised DL-based methods.

Interestingly, there is a positive correlation between shared and state-specific percentages or coverages for the non-DL-based methods (R^2^: 0.92/0.84), while this correlation is negative for supervised DL-based methods (R^2^: 0.96/0.80) (Fig. 2). Similarly, the correlation between precision and the percentage of state-specific contacts is positive for non-DL-based methods (R^2^: 0.95), but negative for supervised DL-based methods (R^2^: 0.91) (Fig. S8). The positive linear relationship indicates that non-DL-based methods simultaneously enhance the prediction accuracy of both shared and state-specific contacts, which collectively contribute to improvements in precision. This can be understood as these methods primarily improve by introducing better strategies to separate direct from indirect couplings. On the other hand, the negative linear relationship suggests that supervised DL-based methods considerably enhanced shared contact prediction while slightly impairing state-specific contact prediction. Despite this, the overall precision still improves by a large margin. Given that the precisions of the contacts maps predicted by supervised DL-based methods improves by a large margin and exceeds 0.9, there are few falsely predicted contacts. This suggests that these supervised DL-based methods prioritized shared contact over state-specific to achieve higher overall precision.

The ESM-1b model parallels the trajectory of non-DL-based methods as seen in Fig. 2. It consistently outperforms CCMpred by 12.3% and 22.4% in percentage, and 15.2% and 28.0% in coverage for shared and state-specific contacts, respectively. This demonstrates the potential advantages of protein language models trained solely on sequence data, thereby avoiding potential biases in current experimental protein structure databases. It would be intriguing to observe whether the continuous improvement of protein language models can concurrently increase the accuracy of shared and state-specific contact prediction, and the IOMemP dataset could serve as a benchmark set for such studies.

### 3.3 Consistency in preferable states across PSP methods

After profiling the performance of PSP methods on shared and state-specific contacts, we analyze their preferences for the IF or the OF conformational states. The “preferable state” is defined to be the state for which a method’s two contact maps consistently yield a higher percentage than the other for the top L long-rage contacts, as introduced in section 2.4.2. We recognized the preferable state for each protein using 11 PSP methods and summarized the results in Fig. 3 (Table S3). The MI method has been excluded from the analysis due to its poor performance on state-specific states.

**Figure 3:**
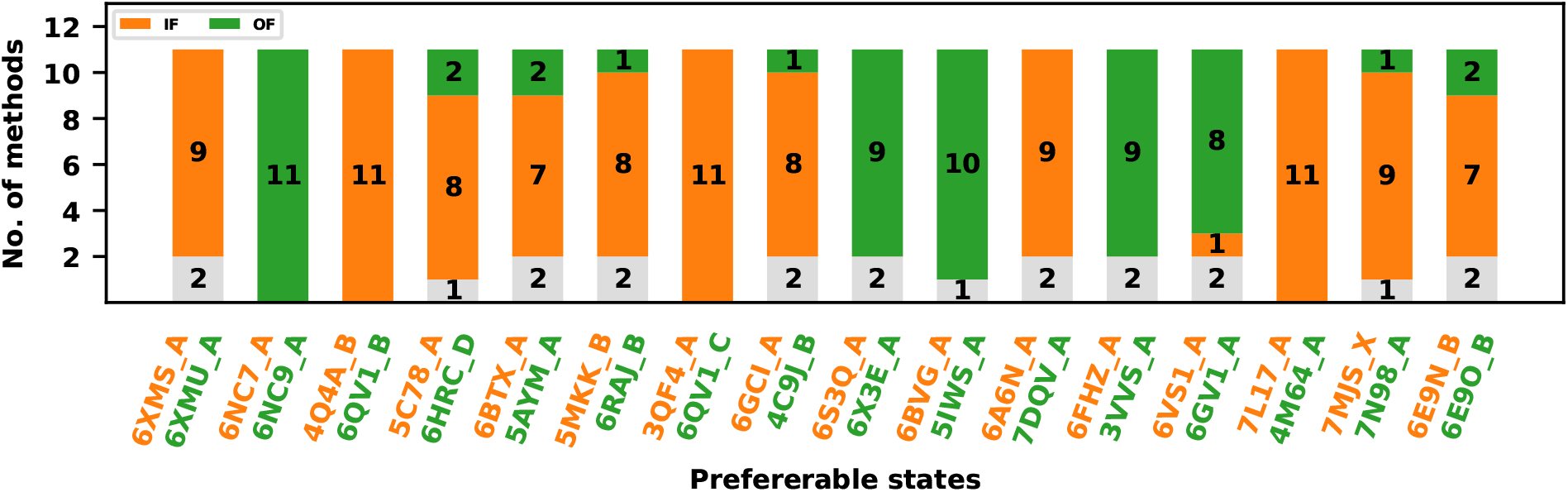
The number of methods that prefer IF (orange) or OF (green) state, or display no state preference (grey).

Our analysis revealed a remarkably consistent preference for specific states across various PSP methods. Among the 16 proteins examined, 11 proteins’ IF states and 5 proteins’ OF states were favored by a substantial majority of PSP methods, while the opposite states were preferred by at most one or two methods. The high degree of agreement in preferable states suggests that the common coevolution input primarily determines which state is perferred. Nevertheless, some methods still exhibited preference for the opposite states, with the specific methods differing for each protein (Fig. S9, S10, and Table S4). The preference of some methods for IF states and others for OF states implies that the input coevolution incorporates information from both states, albeit with a certain degree of bias. The variability suggests that the specific architecture of a given method could potentially influence how the coevolution information is processed for certian sequences, although the effect is not tunable.

We note that for some proteins there were 1 or 2 methods that demonstrated no discernible preference for either state (grey in Fig. 3). Take the protein mdfA for example, trRosetta showed a preference for the IF state (6VS1 A) when its input MSA was generated using the OF structure’s sequence, and for the OF state (6GV1 A) when derived from the IF structure’s sequence (see Fig. S10). The two sequences exhibit very high similarity (91.6% sequence identity) and, importantly, neither 6GV1 A nor 6VS1 A was part of the training data (as of May 1st, 2018) for trRosetta. For this particular protein, it seems that trRosetta’s prediction might fluctuate between the two conformational states given the slight difference in the input of MSAs and sequences.

We then quantified the degree of state preference using either percentage (*d*_percentage_) or coverage (*d*_coverage_) as defined in Eq. 7 and 8, and averaged over the 16 proteins. In general, the *d* values exceed 0.6 and show an increasing trend with more recent PSP methods, consistent with our discussions in previous subsections (Fig. 4 and S11). The unsupervised ESM-1b model emerges again as a outliner that has the lowest preference degrees (*d*_percentage_ = 0.686 and *d*_coverage_ = 0.622), indicating a somehow balanced representation of both IF and OF states. All supervised DL-based methods demonstrate higher preference degrees than ESM-1b, with AlphaFold having highest preference degrees (*d*_percentage_ = 0.919 and *d*_coverage_ = 0.913). This observation reinforces the hypothesis that supervision with experimental structures could be a key factor contributing to the high degree of state preference. The preference computed from contacts was verified by the predicted 3D structures for AlphaFold (Fig. S12). For instance, the two structures predicted for mdfA are close to the preferable IF state, with Δ*D* both less than 0.60 Å (Fig. S10 and S13).

**Figure 4:**
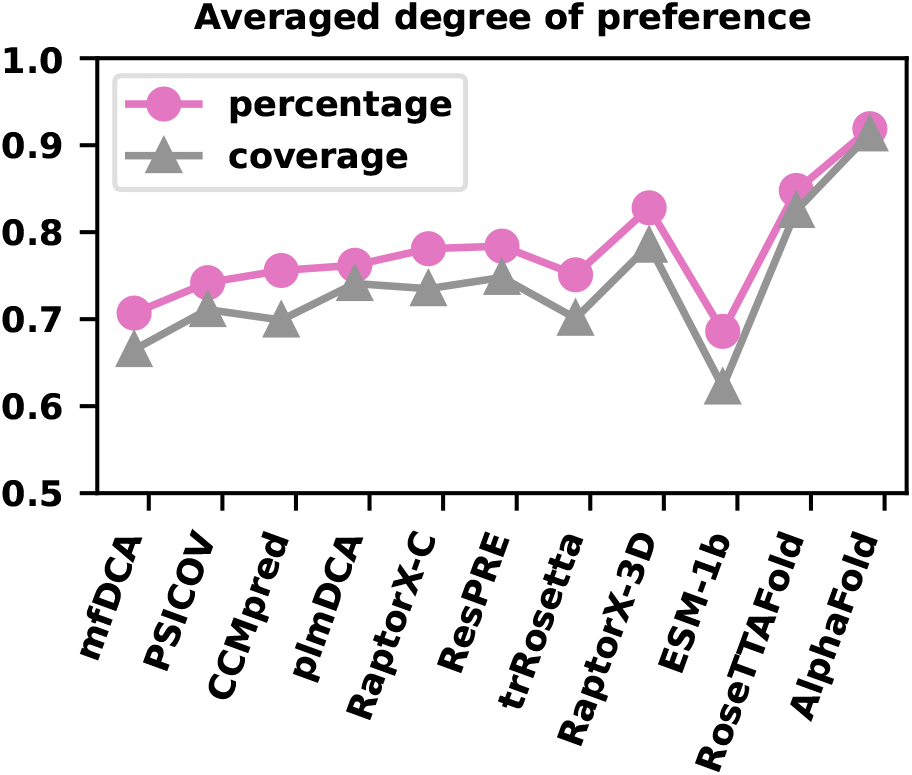
The averaged preference degree *d*_percentage_ (pink) and *d*_coverage_ (gray) for all methods except MI.

Our analysis revealed a predominant preference for the inward-facing conformations, with 11 out of 16 predictions favoring this state. We note that the majority of proteins (12/16) in the IOMemP dataset are involved in substrate efflux while a minority (5/16) involved in influx (Table 1, with AAC3 facilitates both efflux and influx). For transporters that function for substrate efflux, a more stable IF state can be advantageous in the initial substrate binding process and thus exert corresponding evolution pressures that can be manifested as a coevolutionary bias towards the IF states [43, 44]. However, no significant correlation between the preference of PSP predictions and the direction of substrate transport was observed. Among the 12 efflux-related proteins, nine showed a preference for the IF state, while three favored the OF state. Conversely, among the five influx-related proteins, three showed preference for the IF state and two for the OF state. This is likely due to the limited size of the IOMemP dataset and might warrant further investigations.

### 3.4 Limited robustness of ACP methods

Very recently several multiple-state prediction methods have been proposed to predict alternative conformations, and the IOMemP dataset can provide an unbiased benchmark for these ACP approaches. Here we test two popular methods proposed by del Alamo et al [6] and Wayment-Steele et al [7], which use different strategies to extract multiple MSA subsets and subsequently perform individual AlphaFold predictions for each subset. The first method, AF-depth, performs 50 rounds of stochastic sampling from the original MSAs at various MSA depths (8, 16, 32, 64, 128), yielding a total of 250 MSA subsets. Each of these subsets is subsequently used to generate an AlphaFold prediction. In addition, AF-depth carries out predictions using the default AlphaFold MSA depth of 5,120, generating five models. As a result, a conformational ensemble consisting of 255 models is produced for a given protein. The other method, AF-cluster, uses DBSCAN [45] to cluster the MSAs by sequence similarity. Each of these clusters in the MSA space is considered an MSA subset and used independently as an input for AlphaFold predictions. We set the maximum distance (*ϵ*) to 30 to ensure sufficient scanning in DBSCAN.

Having obtained multiple structures from both AF-depth and AF-cluster methods, we scrutinized the presence of two categories of structures. One category closely resembles the IF state (Δ*D*^IF^ *<* Δ*D*_max_*/*2) while differs significantly from the OF state (Δ*D*^OF^ *>* Δ*D*_max_*/*2), and the other category closely matches the OF state while contrasts with the IF state. Finding structures in both categories among all the predicted structures is considered a successful sampling of both the IF and OF states. Otherwise, it is deemed unsuccessful.

Within the IOMemP’s set of 16 proteins, AF-depth successfully predicted both IF and OF states for 7 proteins while AF-cluster achieved successful ACP for 3 proteins. Interestingly, all cases successfully predicted by AF-cluster (AAC3, PF0708, and A5U30 003247) were also successfully predicted by AF-depth. Several exemplary predictions are displayed in Fig. 5. A successful case of ACP would involve identifying conformations that closely match the experimental IF structure and the OF structure (states in the top left and the bottom right regions in Fig. 5), along with a sequence of conformations that transition between these two states. This is exemplified by the AF-depth prediction of SLC1A1, as demonstrated in Fig. 5A. For AAC3 (6GCI A and 4C9J B), both AF-depth and AF-cluster managed to predict alternative conformations, although AF-depth generated more OF state conformations than AF-cluster (see Fig. S14 and 5E). All successful cases are illustrated in Fig. S14 and S15.

**Figure 5:**
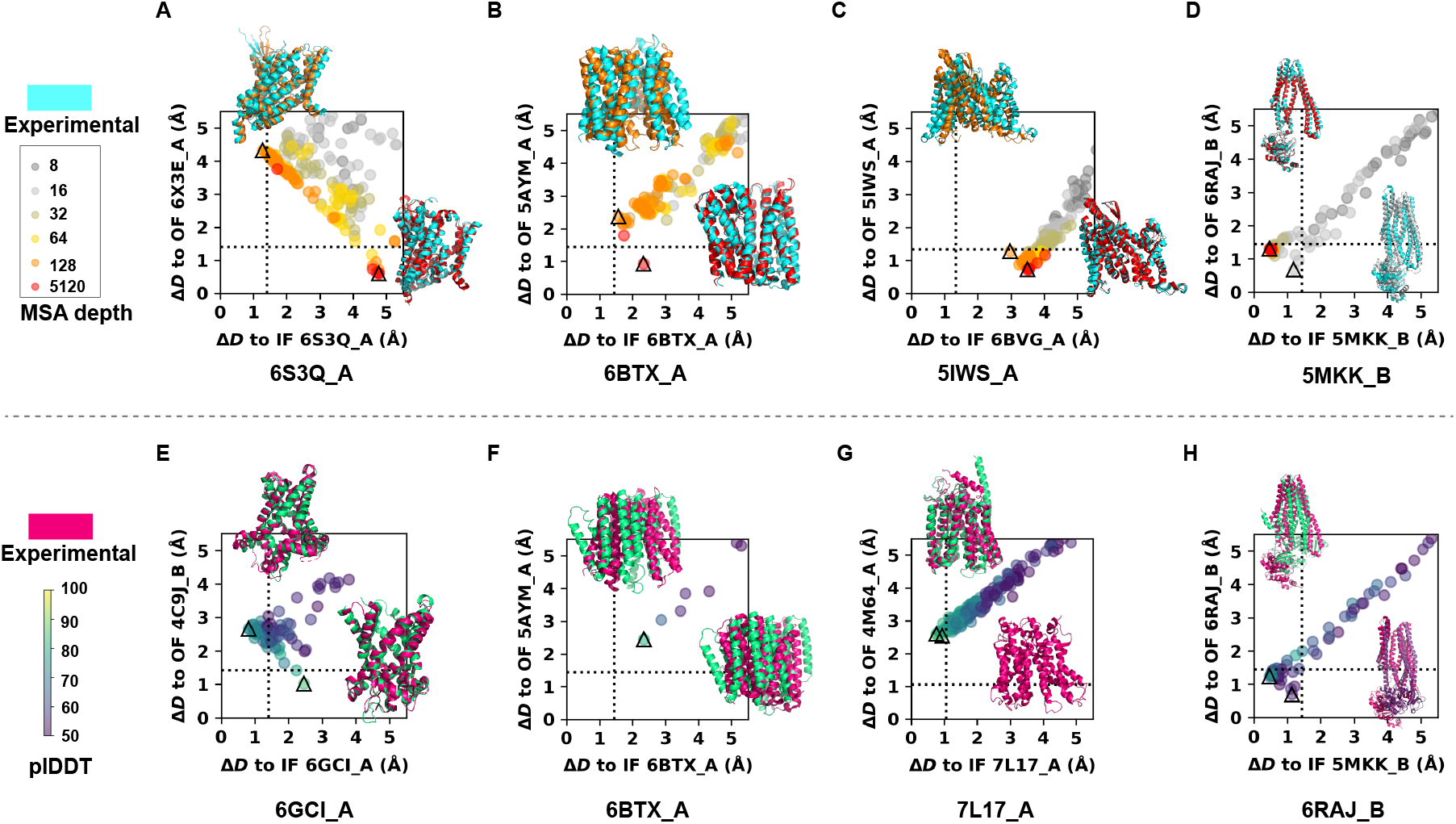
Representative predictions by AF-depth (top panels, A-D) and AF-cluster (bottom panels, E-H). In each panel, the values of Δ*D* of predicted structures with respect to the IF state (x-axis) or to the OF state (y-axis) for a protein are plotted. The target sequences are obtained from either IF or OF structures, and the corresponding PDB and chain IDs are provided. The dotted line in the plot represents Δ*D*_max_*/*2, referring to the deviation between the IF and the OF states. The triangles highlight predicted structures that are closest to either the IF or OF states; these structures are aligned to their closest state (IF or OF) and depicted in cartoon mode (IF: upper left, OF: bottom right). (A-D) Predictions for four proteins by AF-depth. The scatter points and the predicted structures in cartoon mode are colored according to the corresponding MSA depth. Experimental IF and OF structures are shown in cyan. (E-H) Predictions for four proteins by AF-cluster. The scatter points and the predicted structures in cartoon mode are colored according to the corresponding pLDDT values. Only predicted structures with pLDDT values equal to or greater than 50 are considered. Experimental IF and OF structures are shown in magentas.

Unsuccessful predictions present in three distinct situations. The first situation arises when all predicted structures are far from either of the two states, as demonstrated by AF-cluster’s prediction for slc39 (Fig. 5F). We note that a similar outcome was produced with AF-depth when the MSA depth was reduced to less than 128, although the OF state can be recovered using the “complete” MSA (Fig. 5B). This implies that MSA subsets, derived through either method, may eliminate too much information to ensure reliable prediction for slc39.

The second situation involves cases where all predicted structures deviate significantly from one state, with some appearing closer to the other state (Fig. 5C and 5G). This corresponds to the case when stochastic sampling or clustering of MSAs cannot consistently steer AlphaFold toward predicting an alternative state. Possible reasons include that the coevolution information does not contain enough details of the alternative state, or the information of the alternative state is heavily diluted by the other state. The third scenario occurs when some predicted structures align closely with both alternative states judged by Δ*D*, as seen with AF-depth’s and AF-cluster’s predictions for TT C0976 (Fig. 5D and 5H). In the IOMemP dataset, this situation only arose in ABC transporters, whose TM helices rearrangements are not adequately captured by Δ*D*. Comparison with the IF and OF structures suggests that the ACP fails to recapture the experimental conformations.

In summary, the robustness of the two benchmarked ACP methods was found to be limited. For each sequence, AF-depth generated 255 conformations while AF-cluster produced an average of 91 conformations (Table S5. However, in only 44% of cases (7 out of 16) for AF-depth and 19% (3 out of 16) for AF-cluster were both IF and OF structures included among these predicted conformations. Compared to the stochastic MSA subsampling method of AF-depth, AF-cluster presents a more deterministic approach of dividing the “complete” MSA into subsets. The stochastic subsampling, while generating MSA subsets similar to those obtained through clustering, also likely generates a broader array of diverse MSA subsets, some of which may bias towards the alternative state, thus achieving double the success rate for the IOMemP dataset. Exploring more precise and controllable methods of MSA subsampling, such as descent gradient in the MSA space [46], would be beneficial for constructing better ACP methods. The dataset presented in this work could serve as a valuable resource for benchmarking and improving these methods.

### 3.5 Composite confidence scores suggest similar energies between alternative states

Seeking to understand the inherent preferences of the AlphaFold model for alternative protein states, we calculated the composite confidence scores for both IF and OF structures with AlphaFold [32]. The primary hypothesis is that AlphaFold has potentially already learned an approximate energy function capable of estimating the folding free energy of various structural conformations, and the coevolution data (MSA) are used to address the searching problem on this folding energy funnel [47] by providing reasonable initial guesses [32]. This also explains partly the remarkable consistency in preferable states across different PSP methods as discussed in Section 3.3, given that similar initial guesses were generated through unified MSA inputs.

To compute the *s* score, we provided AlphaFold with the target sequence and either an IF or OF structure as the template, eschewing the use of coevolution data. Generally, the output structure from AlphaFold closely resembles the input template structure (TM-score *>* 0.8) and the states adopted are consistent (Fig. S16), with a notable exception of melB. Most importantly, the *s* scores were found to be similar between the two states (Fig. 6). This observation suggests that the AlphaFold energy model, when assessed via the composite confidence score, does not exhibit bias towards a specific alternative state. It further infers that the energy function learned by AlphaFold recognizes both the IF and OF states as local minima in the protein folding energy landscape. Moreover, regardless of whether the target sequences for AlphaFold are derived from IF or OF structures, the composite confidence scores for IF and OF structures are similar for each protein (Fig. 6). This implies that the AlphaFold energy model is largely insensitive to the minor sequence differences between IF and OF structures.

**Figure 6:**
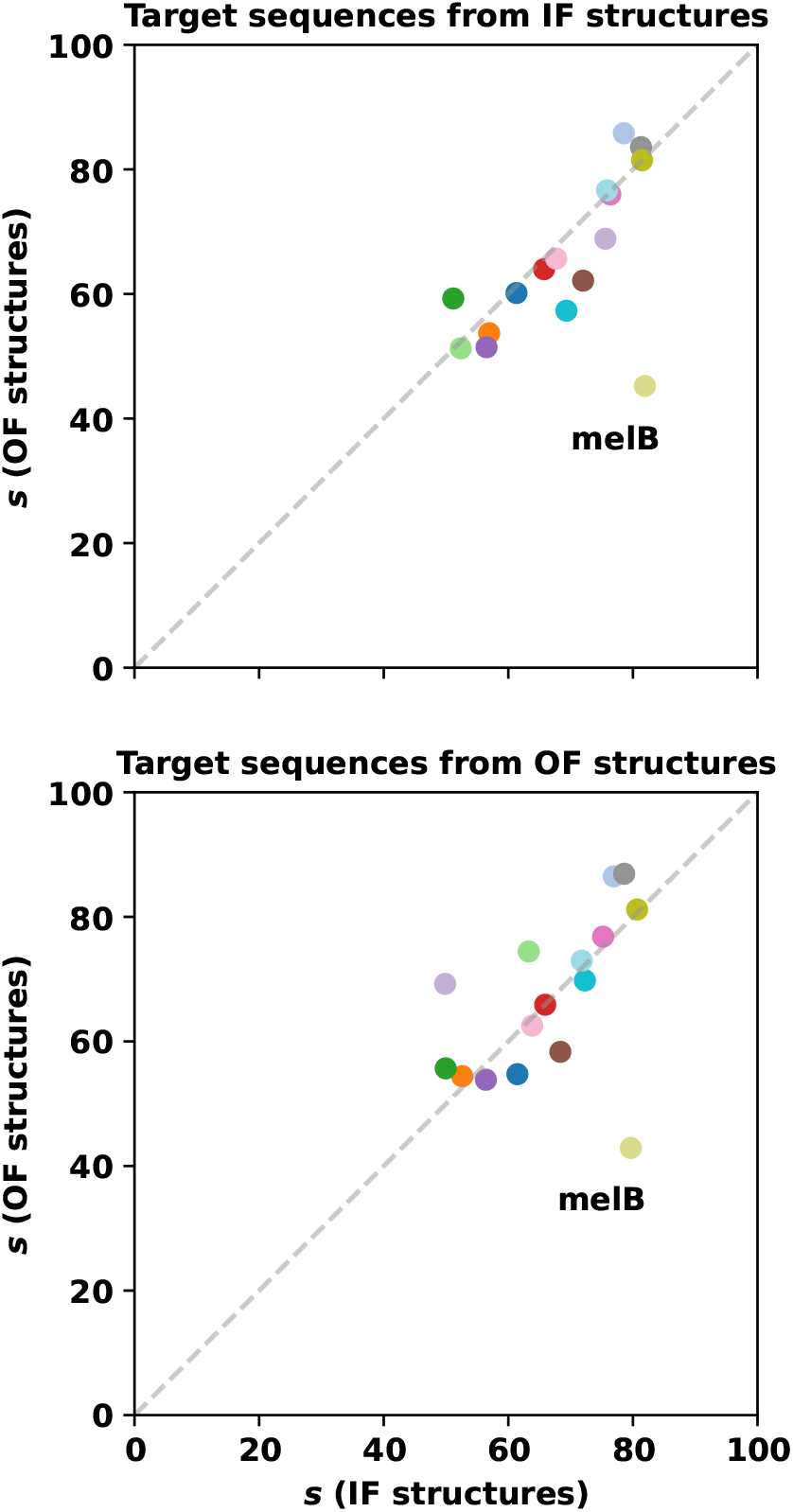
The composite confidence score (*s*) of IF (x-axis) and OF structures (y-axis). Results using target sequences derived from IF structures are shown in the top panel, while those from OF structures are displayed in the bottom panel. Each protein is represented by a unique color. The outliner melB is marked.

Among the 16 proteins in IOMemP, melB represents an exception as its *s* score for the experimental OF structure is significantly lower than that of the IF structure (Fig. 6). This indicates the corresponding OF structure was perceived to be energetically less favorable compared to the IF structure by the energy evaluation module in AlphaFold. Given the limited size of the dataset, it’s uncertain whether this is a common occurrence. We do note that the OF structure (4M64, chain A) was identified as an outward partially occluded conformation [48], so it is possible that it is an intermediate state during transport and a fully OF conformation could potentially yield a higher *s* score.

### 3.6 AlphaFold bias IF when both experimental structures were provided as templates

During the calculation of composite confidence score, one experimental structure (either IF or OF) was provided as the template and no MSA was used. This contrasts with the standard AlphaFold prediction in which the “complete” MSAs and no template were utilized, as carried out in sections 3.1, 3.2 and 3.3. In this analysis, we further investigate the influence of MSAs and template structures (IF and OF) on the state preference manifested by AlphaFold. To this end, we performed AlphaFold predictions using both the experimental IF and OF structures simultaneously as the input template. This approach resulted in three sets of predictions: one using only the template as input (Fig. S17), another using only the MSAs (Fig. S18), and a third using both the MSAs and the template (Fig. S19). It is important to note that these predictions employed AlphaFold model 1 with a single recycling iteration and were repeated 25 times for each input configuration, diverging from the default AlphaFold settings using 5 models with 3 recycling iterations.

AlphaFold predicted the same state between the first two sets of predictions (one using only template and another using only MSAs) for most cases (13 out of 16) (Fig. S17 and S18). More notably, in all these cases the IF conformational states were predicted. This tendency towards IF states might be attributed to AlphaFold’s learning that IF states are more evolutionarily favorable [43, 44]. Despite recognizing both IF and OF states as local energy minima based on the composite confidence score, AlphaFold displays a general preference for the IF states. Once “complete” MSAs were used as input, the predictions from AlphaFold were largely identical, regardless of whether IF and OF structures were provided as templates (Fig. S18 and S19). This suggests that coevolution information profoundly directs AlphaFold towards a specific state, a bias that appears unaffected by template inputs. Such an observation thus aligns well with the assessment that MSAs dictate the predicted conformation in ACP.

## 4 Discussion and Conclusions

The Critical Assessment of protein Structure Prediction (CASP) sought to provide targets for predicting “protein conformational ensembles” in CASP15 (2022), which highlighted the importance of evaluating methods for predicting alternative conformations and structural ensembles [49]. In this study, we curated the IOMemP dataset containing both IF and OF structures of 16 membrane proteins, and used it to benchmark 12 PSP methods and 2 multiple-state methods for alternative state predictions. The general goal was to provide theoretical insights into the ACP problem and to facilitate ongoing advancement to address the problem.

Among the 12 PSP methods, the encouraging improvements in contact prediction were primarily driven by higher percentages of accurately predicted shared contacts. While the adoption of DL architectures such as ResNN and Transformer into supervised learning of experimental structures has signficantly enhanced shared contact predictions, the prediction of state-specific contacts seems to have been somewhat compromised. This contrasts with earlier non-DL-based method development where improvements were observed in the prediction of both shared and state-specific contacts. ESM-1b, a protein language model, exhibited a better-balanced performance in predicting IF and OF states, indicating its potential advantages.

An interesting observation is that these PSP methods exhibited highly consistent state preferences, although the degree of preference varies. Further analysis revealed that such preferences were largely determined by the input MSAs, which were unified for different methods in this study. The MSAs, generated by the AlphaFold protocol, are hypothesized to contain nearly “complete” coevolution information. Our study demonstrated that the manner in which MSAs are handled likely introduces a bias toward a specific state, thereby influencing the preference of these methods for a state. When providing both IF and OF structures as input templates, AlphaFold occasionally predicted both conformational states. However, including MSAs in the input led AlphaFold to predict a single state consistently, affirming the bias introduced by MSAs. Most notably, comparison of the composite confidence scores between the experimental IF and OF structures suggested that the energy function learned by AlphaFold indeed estimates similar free energies for these two alternative states. This suggests that conformational searching or sampling is key to solving the ACP problem.

The IOMemP dataset could function as valuable unbiased resource for benchmarking emerging ACP methodologies. Tests with AF-depth and AF-cluster demonstrated a general lack of robustness, yet they revealed succesful prediction in several instances (7 out of 16 for AF-depth and 3 out of 16 for AF-cluster). These findings underscore the potential for further research to uncover novel strategies for manipulating MSAs or refining their corresponding representations, thus overcoming the observed limitations.

When examining the frequency of the predictions of IF and OF states, it is evident that the IF states are often preferred when provided with “complete” MSAs. Similarly, AlphaFold predominantly selects IF states when only IF and OF structures are used as input templates without including MSAs. These observations hint a greater inherent favorability for IF states relative to OF states. This preference could be due to the greater evolutionary stability of IF states compared to OF states, a fact supported by the general tendency of structural biologists to capture IF states for membrane proteins [44].

At the outset of this study, we hypothesized that the IF and OF structures, characterized by a high sequence identity (*>*70%), belonged to the same protein prior to dataset curation. To validate this hypothesis, we assessed the performance of the 12 PSP methods across different sequences of a target protein. We compared the percentages of their predicted contacts (*p*_sh_, *p*_in_, and *p*_out_) as well as the differences (*p*_in_ − *p*_out_) between sequences of IF and OF structures, and found them to be highly similar (Table S6). Energy estimatation, using the *s* scores, also indicated a close similarity between the sequences of IF and OF structures. Moreover, AlphaFold predictions showed no significant differences between sequences of IF and OF structures when various inputs were used (Fig. S16-S19). These findings collectively suggest that sequence variations did not significantly impact our ACP benchmark. Nevertheless, it may be worth investigating to establish a valid lower bound of sequence identity that allows for accurate alternative conformation predictions.

While we believe that the IOMemP dataset could facilitate the development of ACP methods, we note that it comprises only two types of alternative states for a limit set of single-domain membrane proteins. Expanding this to larger databases, featuring diverse alternative conformations and multiple structural states derived from experimental data, would be beneficial. Moreover, there is a pressing need to establish evaluation metrics capable of assessing the quality of predicted conformations for a broader range of proteins, not only membrane proteins but also globular proteins and multi-domain proteins with diverse structural features. Such efforts will further enrich our understanding of protein conformational ensembles and dynamcis, as well as aid in the development of robust ACP methods that can be broadly applicable across various protein systems.

## Acknowledgements

We thank Drs. Wenyi Zhang, Yingnan Hou, Zilin Song and Xiaoli Lu for the helpful discussions. The work is supported by the “Pioneer” and “Leading Goose” R&D Program of Zhejiang (2023C03109) and the National Natural Science Foundation of China (32171247, 21803057). We thank the Westlake University Supercomputer Center for computational resources and related assistance.

## Author contributions

J.H. conceived the study. T.X. performed the experiments. T.X and J.H. wrote the manuscript together.

## Data and code availability

More detailed information about the IOMemP dataset can be found in Data S1, while the versions and sources of the benchmarked methods are available in Data S2. The code for dataset construction and the benchmarking work has been deposited on GitHub (https://github.com/JingHuangLab/IOMemP). Any additional information required to reanalyze the data reported in the paper can be obtained from the corresponding author upon request.

Data analysis was carried out using Python v3.8, Matplotlib v3.5.0, NumPy v1.19.5, SciPy v1.2.1, pandas v1.3.5, and scikit-learn v1.1.2. TOPCONS2 v1.2 and memembed v1.15 were used for the sequence-based topology prediction and structure-based membrane orientation prediction, respectively. TM-align v20190822 was used for computing TM-scores, GREMLIN CPP v1.0 for computing NEFF, and MMseqs2 for sequence identity. PyMol v2.5.5 was used to create structure visualizations.

## Supplementary Material

**Table S1:** The average number of different contacts with respect to the sequence length (*L*).

| contacts | all-range | long-range | medium-range | short-range |
| --- | --- | --- | --- | --- |
| shared | 4.008 | 3.721 | 0.200 | 0.087 |
| state-specific | 1.501 | 1.435 | 0.043 | 0.023 |
| native (shared + state-specific) | 5.503 | 5.156 | 0.243 | 0.110 |

**Table S2:** The average values of *p*_sh_ + *p*_in_, *p*_sh_ + *p*_out_, and *P* for each method.

| Methods | $p_{\text{sh}} + p_{\text{in}}$ | $p_{\text{sh}} + p_{\text{out}}$ | $P$ |
| --- | --- | --- | --- |
| MI | 0.081 | 0.078 | 0.094 |
| mfDCA | 0.362 | 0.356 | 0.411 |
| PSICOV | 0.311 | 0.294 | 0.347 |
| CCMpred | 0.549 | 0.517 | 0.592 |
| plmDCA | 0.448 | 0.423 | 0.488 |
| RaptorX-Contact | 0.872 | 0.804 | 0.914 |
| ResPRE | 0.862 | 0.793 | 0.904 |
| trRosetta | 0.865 | 0.812 | 0.915 |
| RaptorX-3DModeling | 0.923 | 0.854 | 0.957 |
| ESM-1b | 0.626 | 0.583 | 0.677 |
| RoseTTAFold | 0.923 | 0.881 | 0.961 |
| AlphaFold | 0.953 | 0.931 | 0.998 |

**Table S3:** The number of proteins for which the two predicted contacts from two different sequences both prefer the same states for each method.

| Methods | Number of proteins | Percentage | Proteins not preferred by the method |
| --- | --- | --- | --- |
| MI | 8 (16) | 50% |  |
| mfDCA | 15 (16) | 93.8% | CYME_CMD148C |
| PSICOV | 15 (16) | 93.8% | TT_C0976 |
| CCMpred | 15 (16) | 93.8% | A5U30_003247 |
| plmDCA | 14 (16) | 87.5% | wlaB, AAC3 |
| RaptorX-Contact | 14 (16) | 87.5% | PF0708, MFSD2A |
| ResPRE | 13 (16) | 81.3% | slc39, SLC1A1, mdfA |
| trRosetta | 12 (16) | 75.0% | TT_C0976, PF0708, mdfA, A5U30_003247 |
| RaptorX-3DModeling | 15 (16) | 93.8% | SLC1A1 |
| ESM-1b | 14 (15) | 93.3% | CYME_CMD148C |
| RoseTTAFold | 13 (15) | 86.7% | AAC3, ptsG |
| AlphaFold | 15 (16) | 93.8% | slc39 |

ESM-1b and RoseTTAFold failed in the prediction of SPF1.

**Table S4:** Seven proteins whose IF or OF states are preferred by only 1 or 2 PSP methods.

| IF/OF | PSP methods |
| --- | --- |
| 6HRC_D | mfDCA, AlphaFold |
| 5AYM_A | PSICOV, RoseTTAFold |
| 6RAJ_B | mfDCA |
| 4C9J_B | AlphaFold |
| 6VS1_A | AlphaFold |
| 7N98_A | AlphaFold |
| 6E9O_B | mfDCA, RoseTTAFold |

**Table S5:**
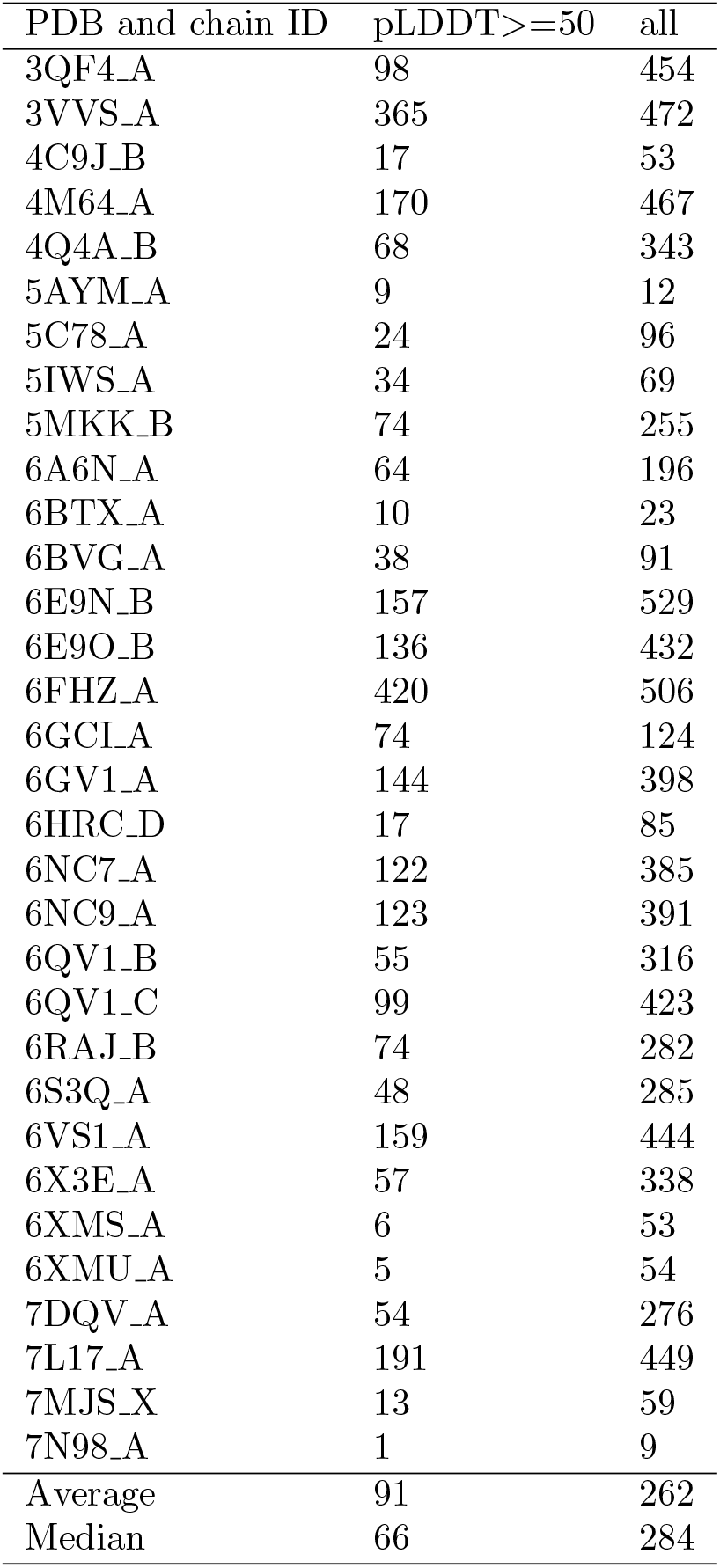
The number of conformations generated by AF-cluster for each target. Only those with pLDDT equal to or larger than 50 were analyzed.

**Table S6:** The fitness (R^2^) of the linear regression (*y* = *x*) for percentages (*p*_sh_, *p*_in_, and *p*_out_) or percentage differences (*p*_in_ − *p*_out_) between IF-derived and OF-derived sequences, illustrating that sequence differences do not significantly impact predictions.

| methods | $R^2$ | | | |
| --- | --- | --- | --- | --- |
| | $p_{\text{sh}}$ | $p_{\text{in}}$ | $p_{\text{out}}$ | $p_{\text{in}} - p_{\text{out}}$ |
| MI | 0.89 | 0.89 | 0.82 | 0.94 |
| mfDCA | 0.96 | 0.93 | 0.78 | 0.83 |
| PSICOV | 0.86 | 0.76 | 0.75 | 0.80 |
| CCMpred | 0.94 | 0.91 | 0.88 | 0.89 |
| plmDCA | 0.97 | 0.89 | 0.87 | 0.87 |
| RaptorX-Contact | 0.92 | 0.87 | 0.73 | 0.85 |
| ResPRE | 0.97 | 0.95 | 0.78 | 0.90 |
| trRosetta | 0.95 | 0.96 | 0.66 | 0.87 |
| RaptorX-3DModeling | 0.86 | 0.94 | 0.82 | 0.92 |
| ESM-1b | 0.96 | 0.97 | 0.39 | 0.91 |
| RoseTTAFold | 0.94 | 0.81 | 0.48 | 0.61 |
| AlphaFold | 0.87 | 0.96 | 0.70 | 0.88 |

**Figure S1:**
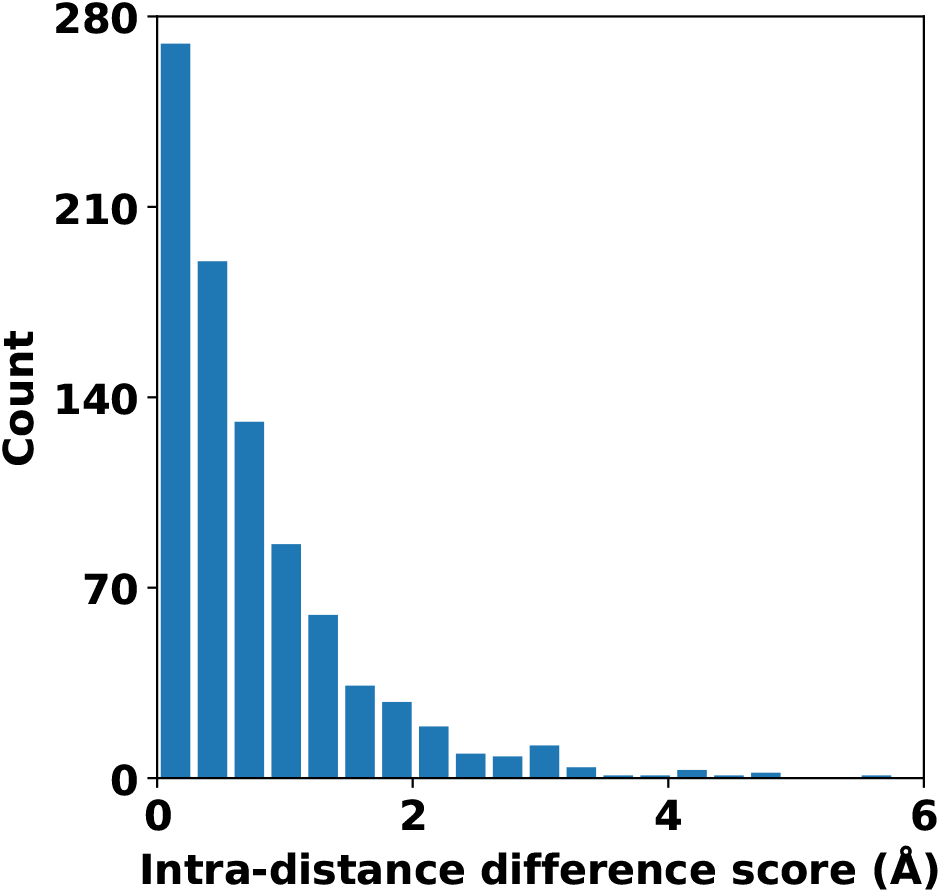
Proteins with large intra-distance differences Δ*D* are extremely scarce.

**Figure S2:**
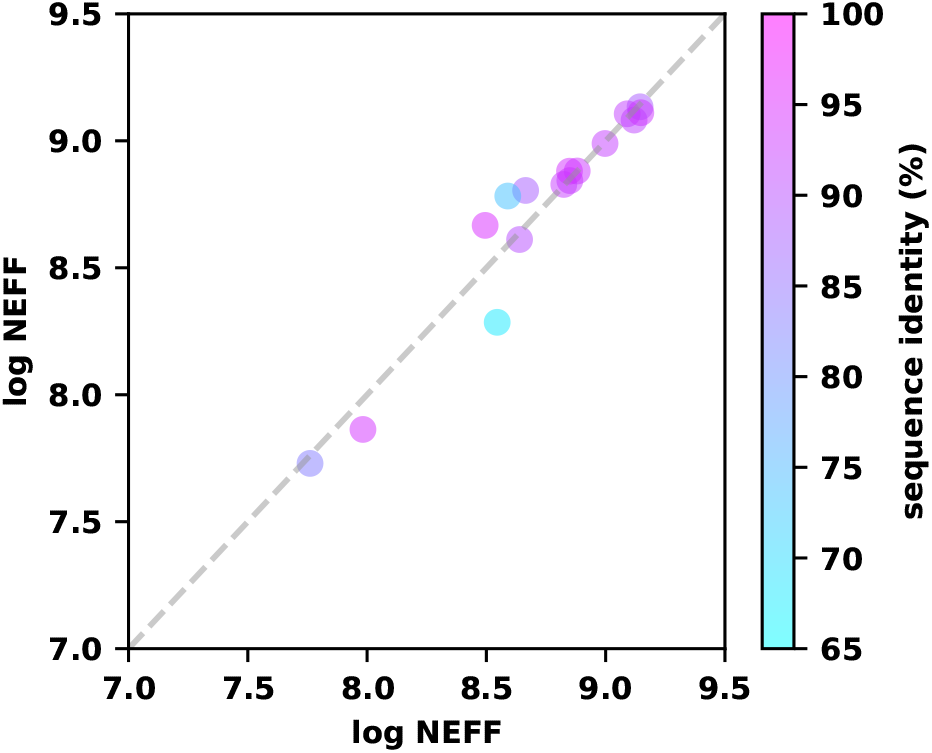
The logarithm values of NEFF for the two sets of MSAs, derived from the sequences of the two states of each protein, show high similarity. The color bar represents the sequence identities between alternative states.

**Figure S3:**
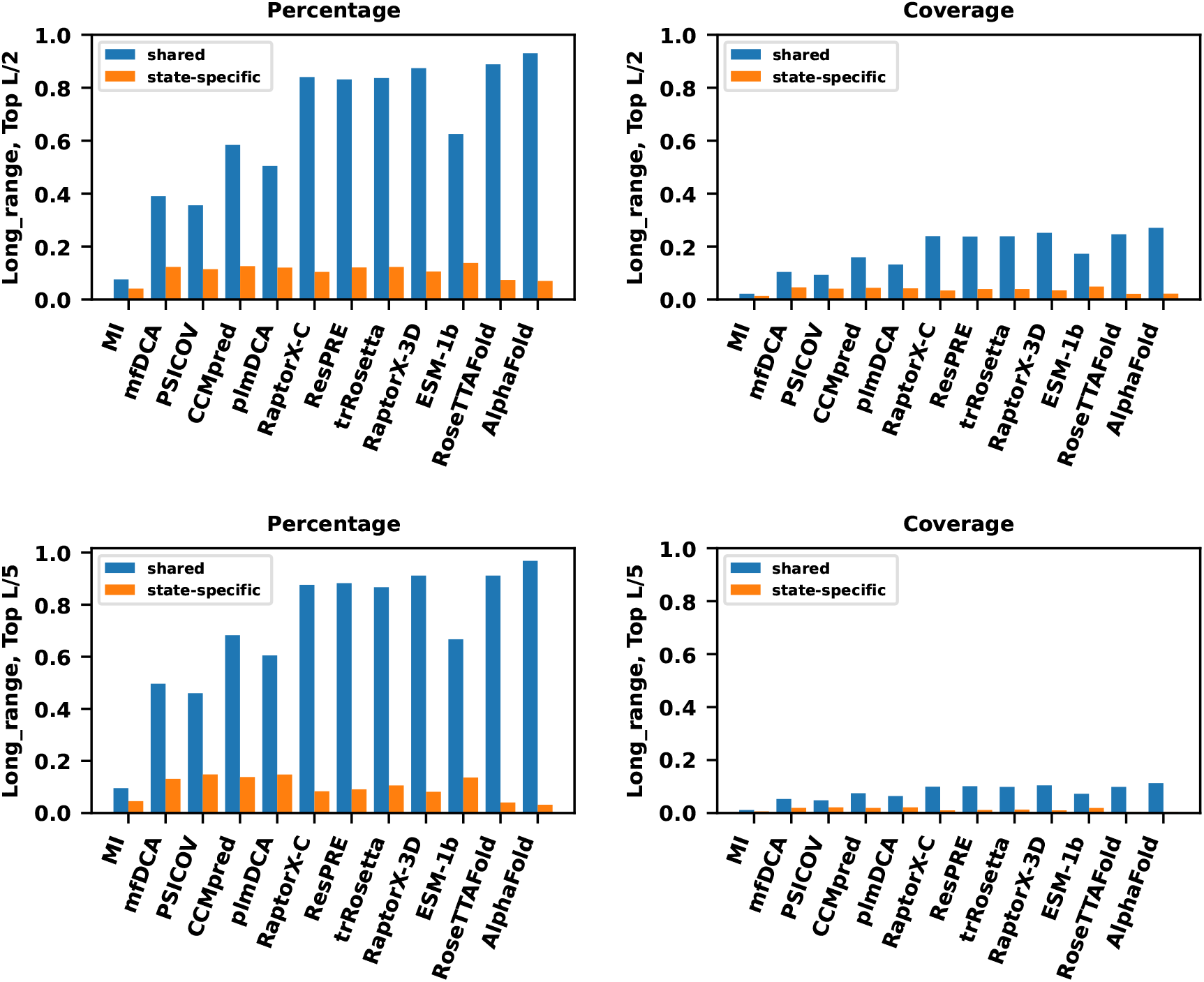
The averaged percentages or coverages of the 12 PSP methods for the top L/2 (or top L/5) long-range contacts.

**Figure S4:**
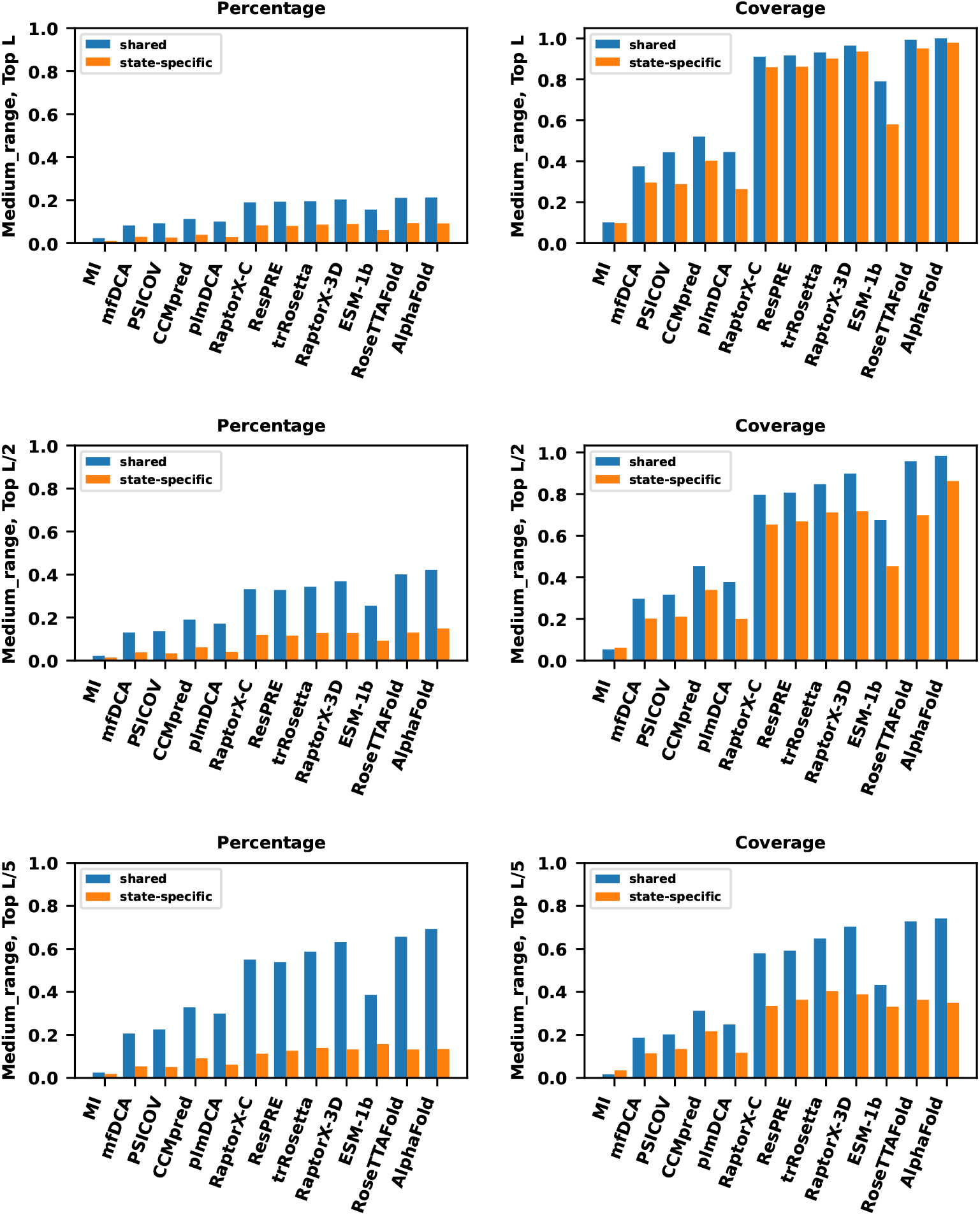
The averaged percentages or coverages of the 12 PSP methods for the top L*/n* medium-range shared (blue) and state-specific (orange) contacts.

**Figure S5:**
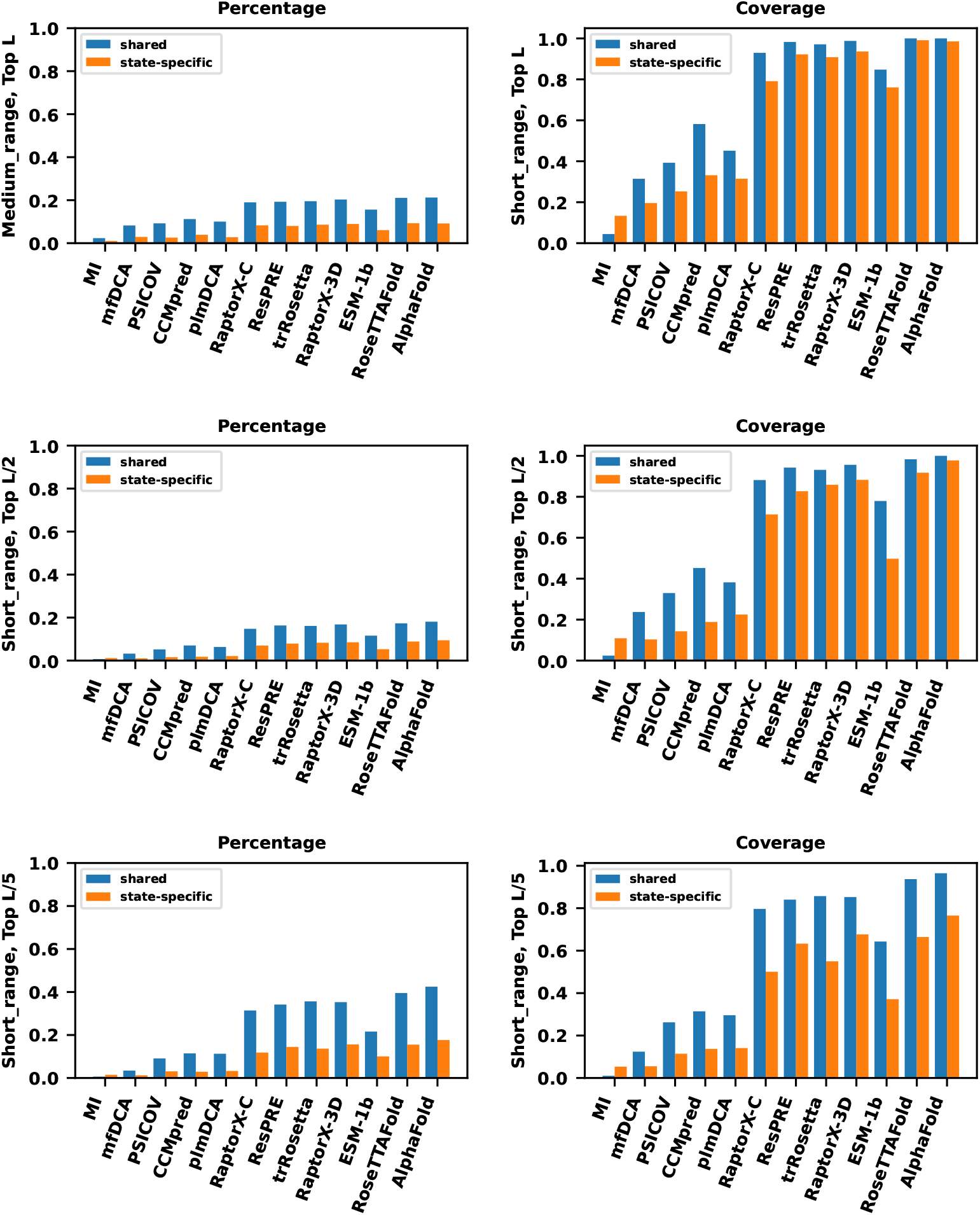
The averaged percentages or coverages of the 12 PSP methods for the top L/*n* short-range shared (blue) and state-specific (orange) contacts.

**Figure S6:**
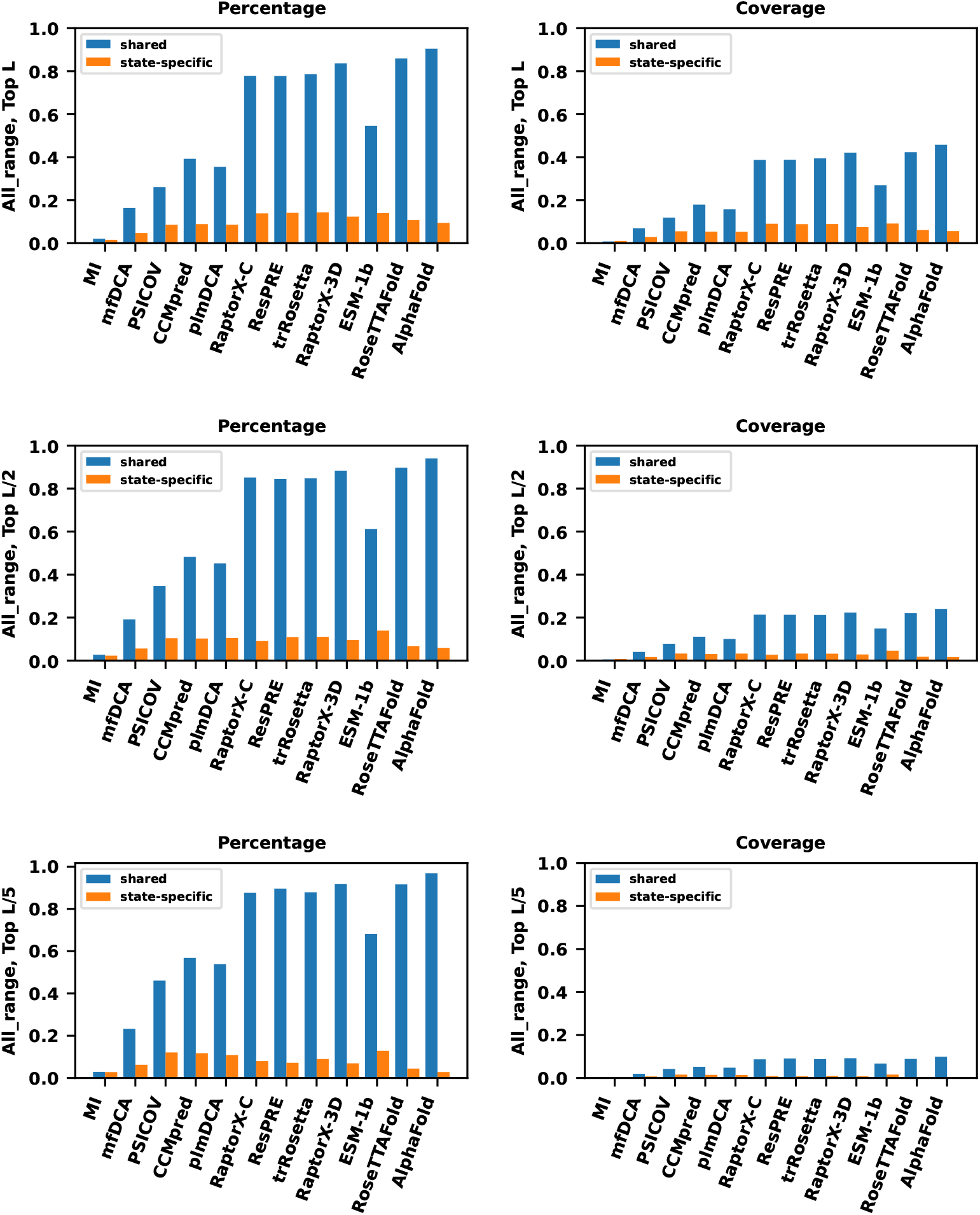
The averaged percentages or coverages of the 12 PSP methods for the top L/*n* all-range shared (blue) and state-specific (orange) contacts.

**Figure S7:**
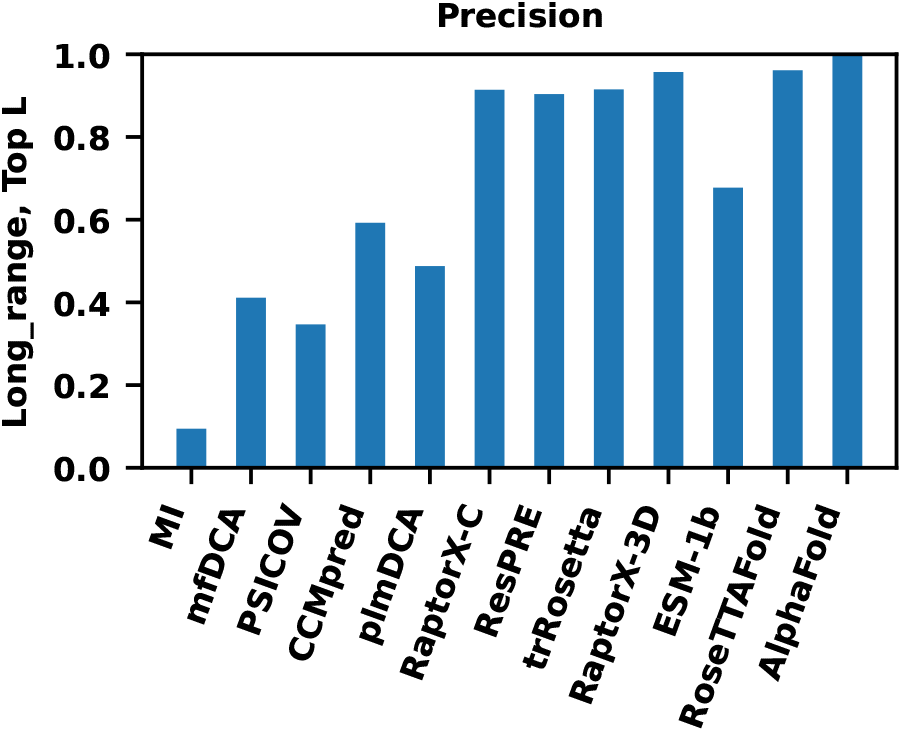
The averaged precision (*P*) of the 12 PSP methods for the top L long-range contacts, where supervised DL-based methods’ precisions exceed 0.9.

**Figure S8:**
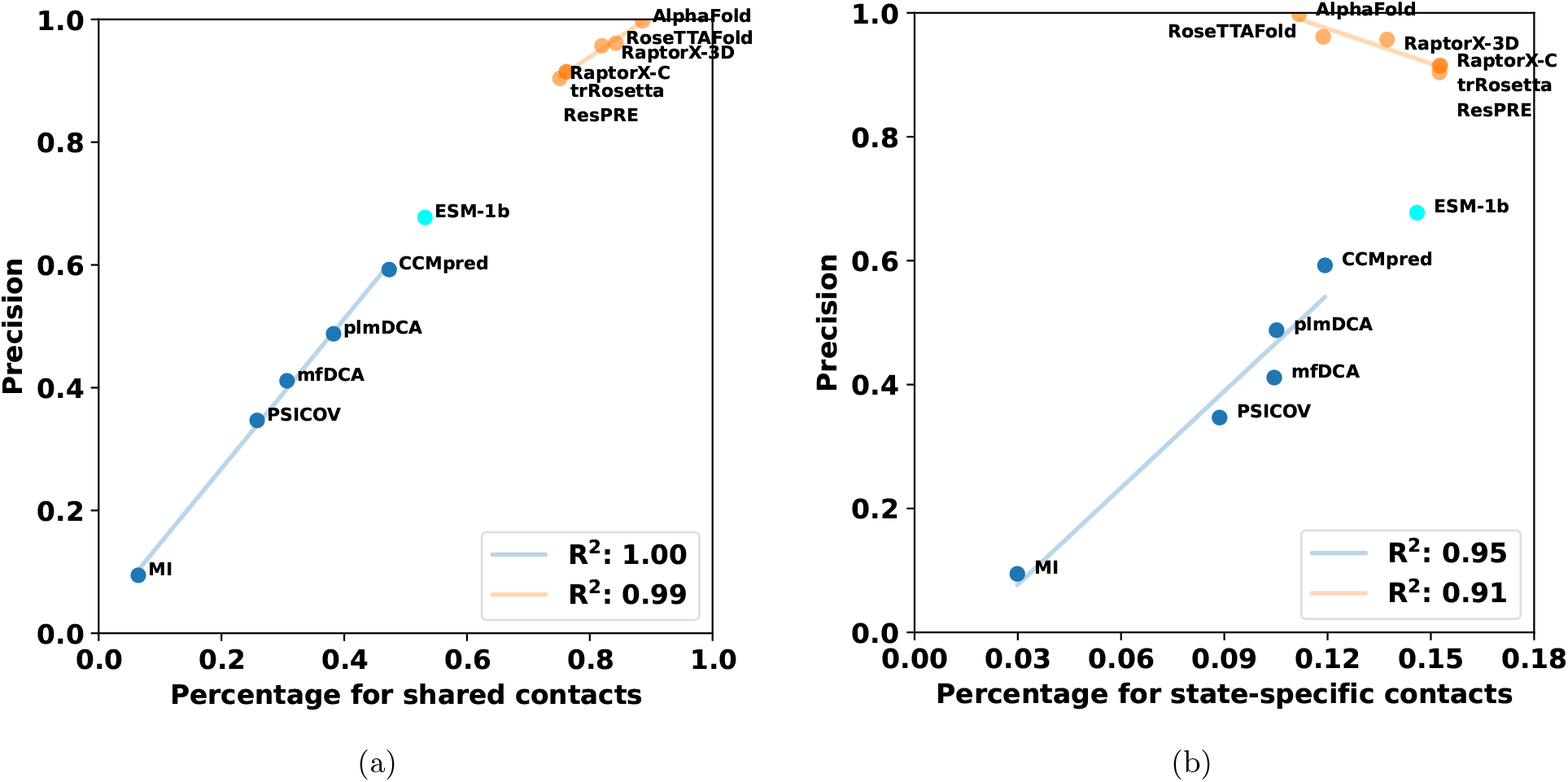
For the top L long-range contacts, the precision versus the percentages for shared (a) and state-specific (b) contacts. Non-DL-based methods exhibit positive correlations in both (a) and (b), while supervised DL-based methods exhibit positive correlations in (a) and negative correlations in (b).

**Figure S9:**
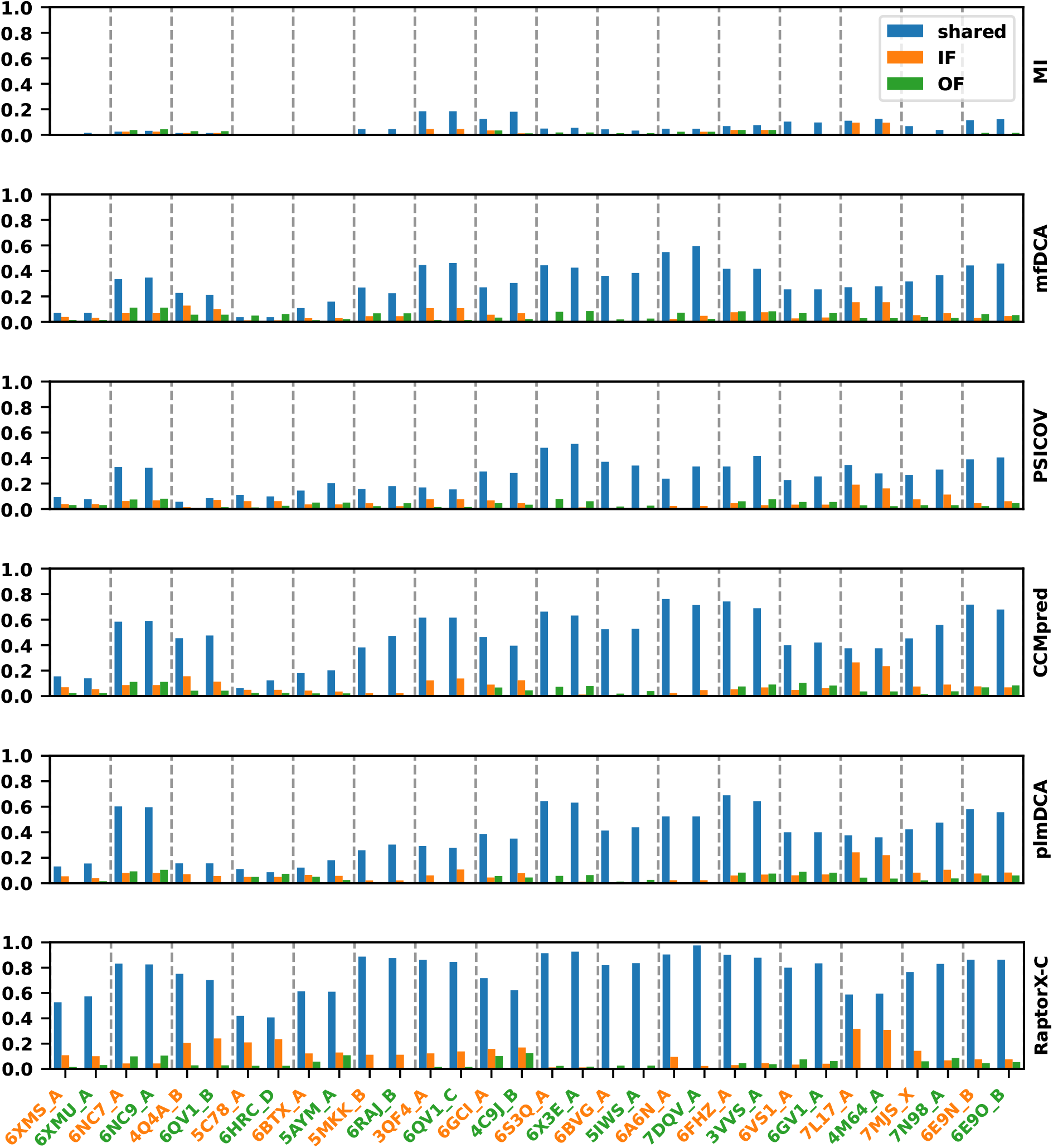
The percentages for the top L long-range contacts for each protein by 6 PSP methods (blue: *p*_sh_; orange: *p*_in_; green: *p*_out_). The gray dashed lines divide the 16 proteins. The PDB IDs are colored according to their states (IF: orange, OF: green), which correspond to the sequences used as target sequences.

**Figure S10:**
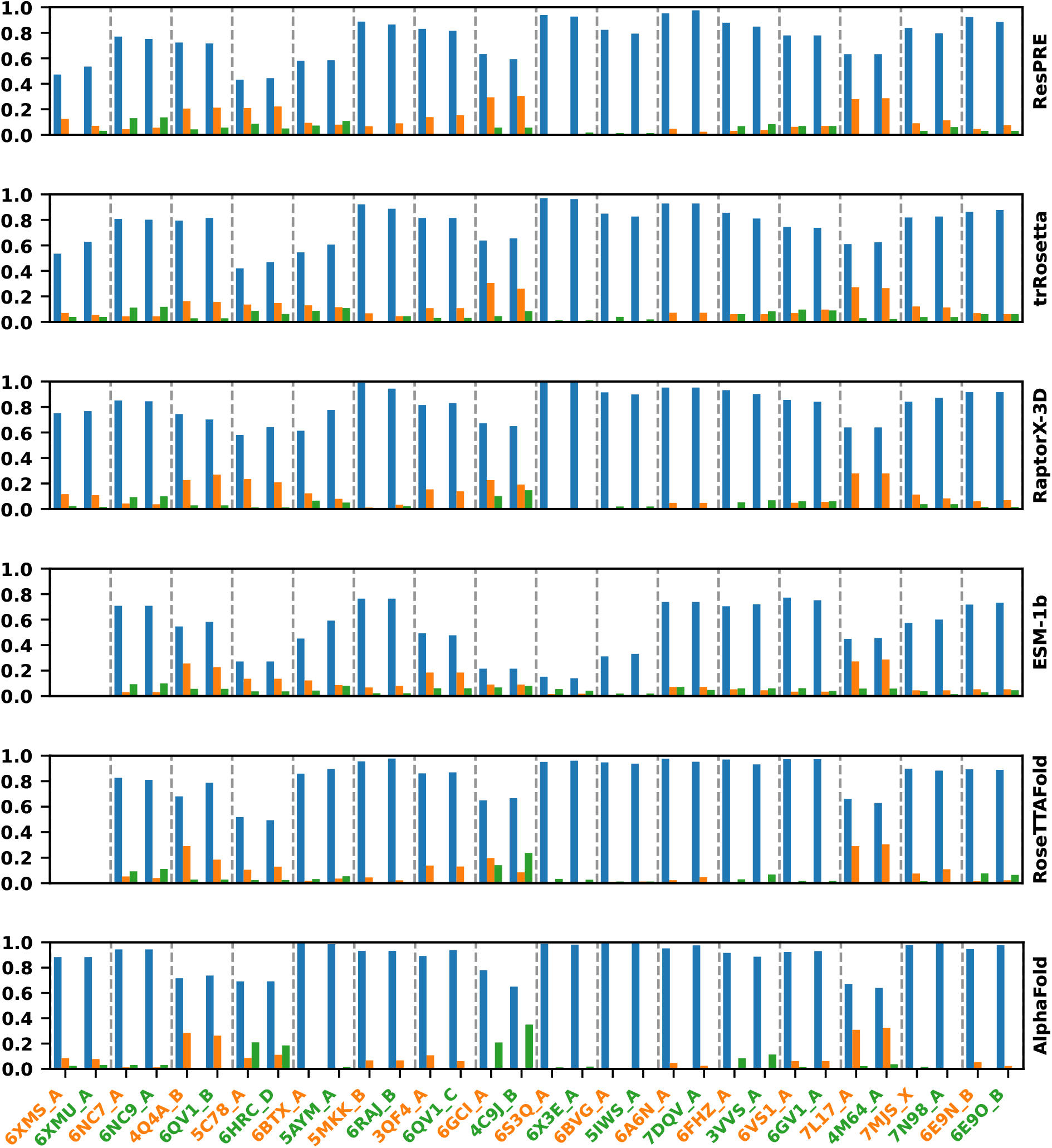
The percentages for the top L long-range contacts for each protein by the remaining 6 PSP methods (blue: *p*_sh_; orange: *p*_in_; green: *p*_out_). The gray dashed lines divide the 16 proteins. The PDB IDs are colored according to their states (IF: orange, OF: green), which correspond to the sequences used as target sequences. The predictions of ESM-1b and RoseTTAFold on 6XMS A and 6XMU A failed due to out of memory.

**Figure S11:**
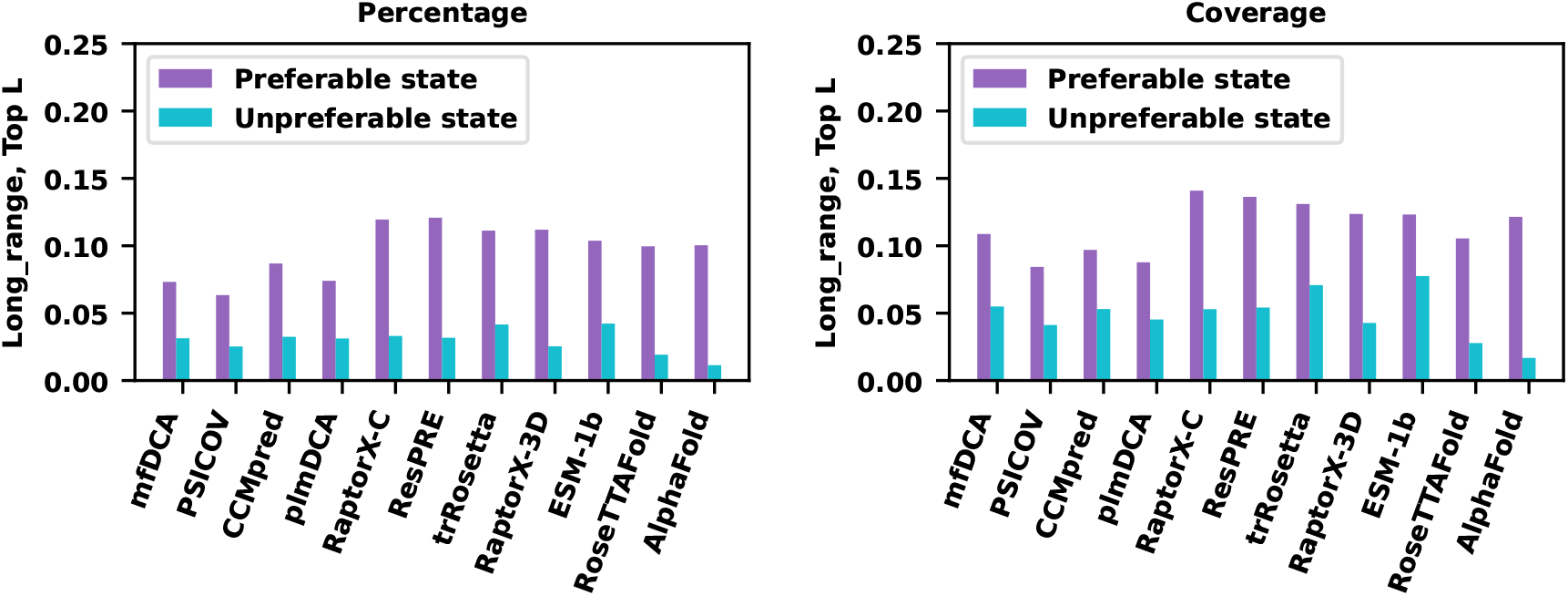
The mean percentages or coverages for the top L long-range contacts for preferable (purple) and unpreferable (blue) states for each method.

**Figure S12:**
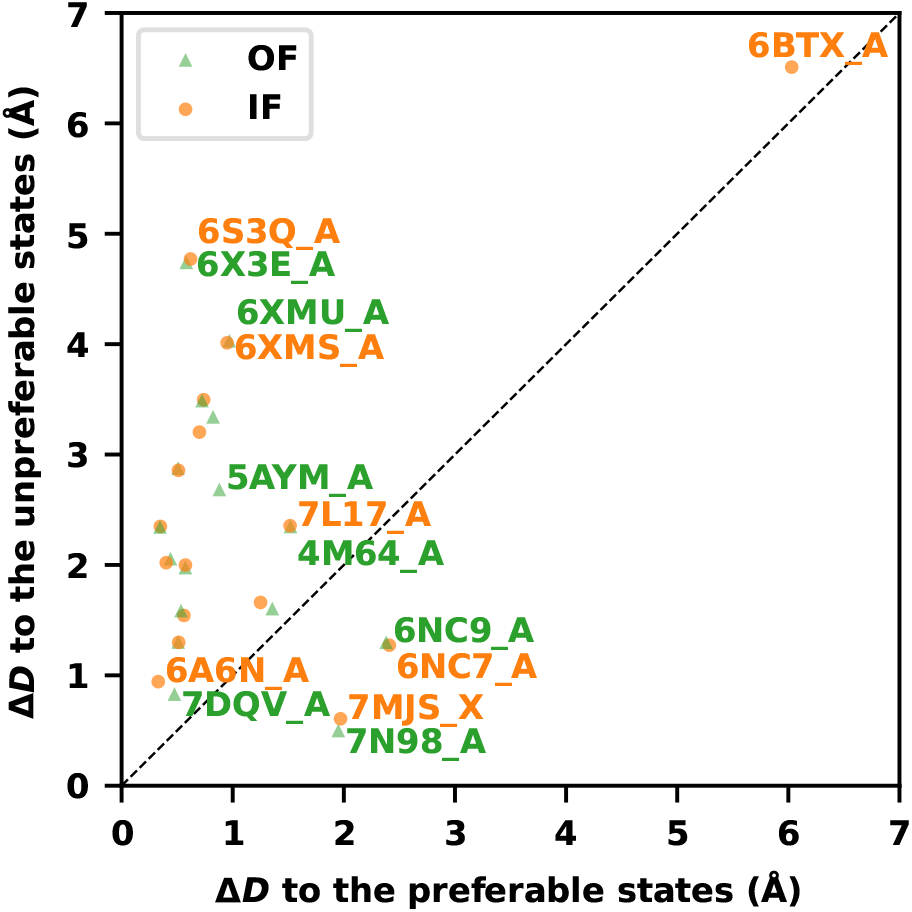
The AlphaFold predicted structures are generally closer to preferable states (smaller Δ*D*) and relatively more distant from unpreferable states (larger Δ*D*), where preferable and unpreferable states are assigned according to the percentage for the top L long-range contacts. Out of the 32 predicted structures, the preferable states of 27 proteins, as implied in AlphaFold contacts, align with the states of AlphaFold predicted structures. The orange points refer to the target sequences derived from IF structures and the green triangles OF structures.

**Figure S13:**
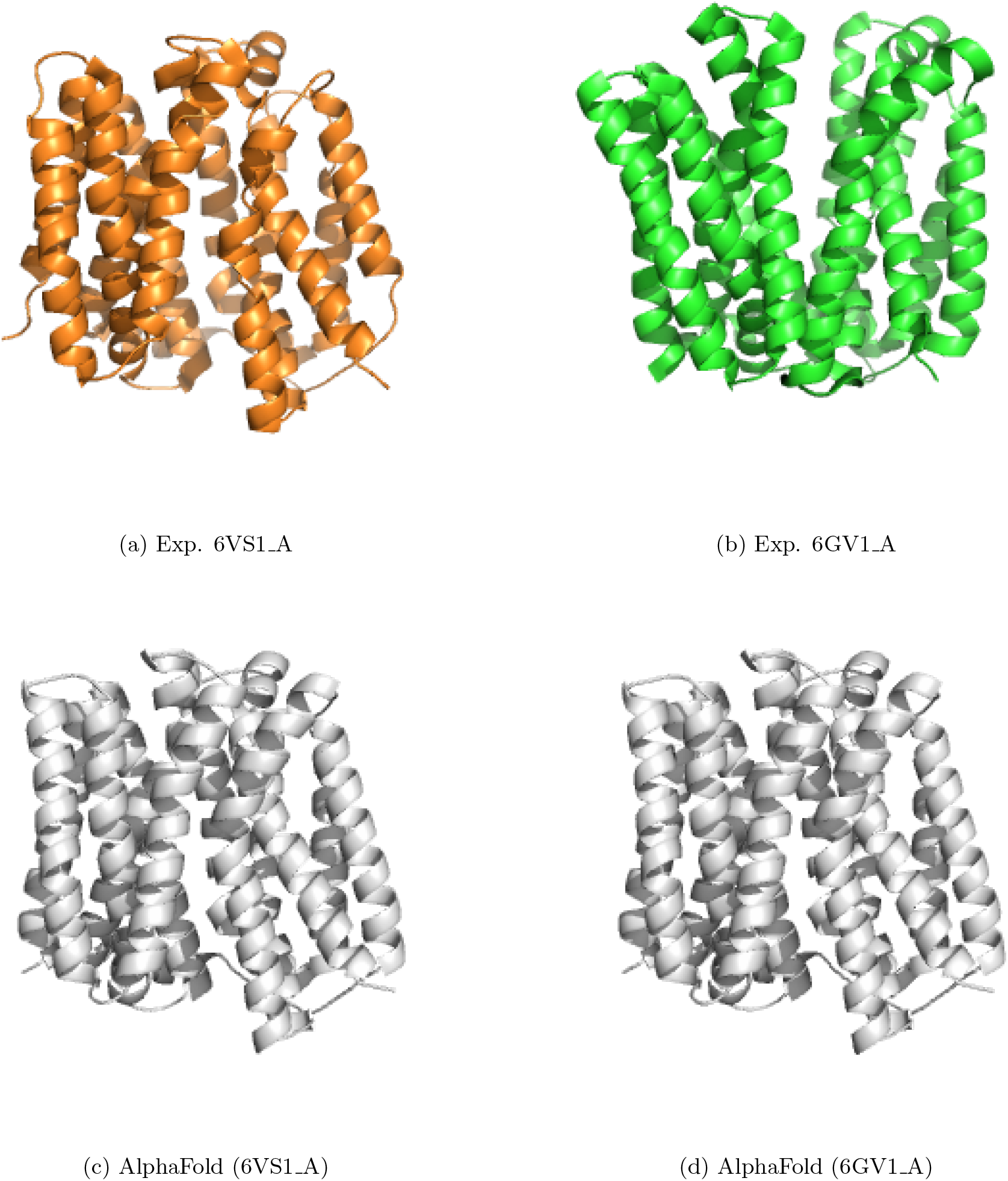
The experimental structures of the Multidrug transporter MdfA in (a) IF and (b) OF structures, along with (c, d) the AlphaFold predicted structures. Both predicted structures align with the IF state, although their respective target sequences are derived separately from 6VS1 A and 6GV1 A.

**Figure S14:**
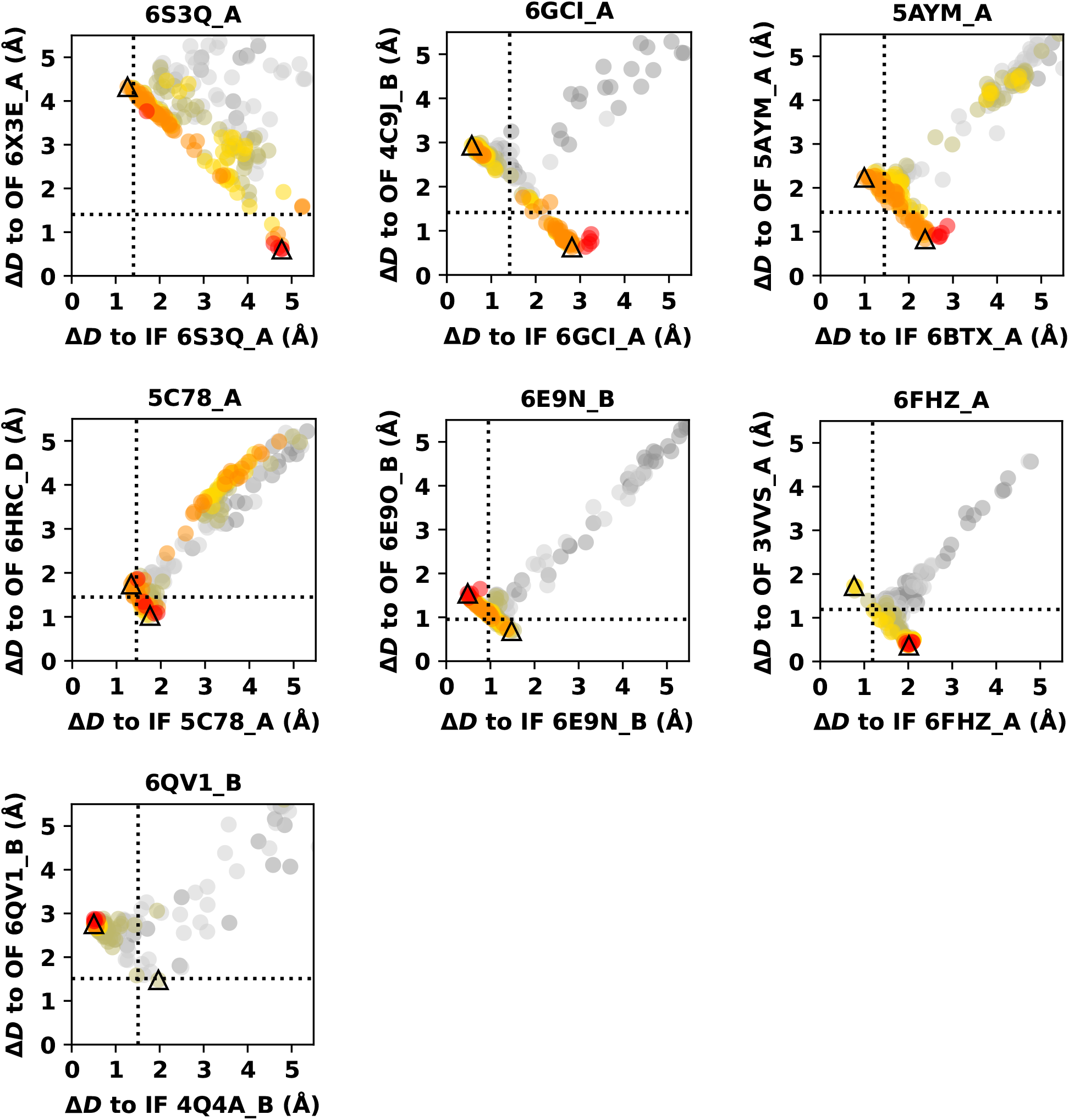
Successful ACP predictions by AF-depth for IOMemP.

**Figure S15:**
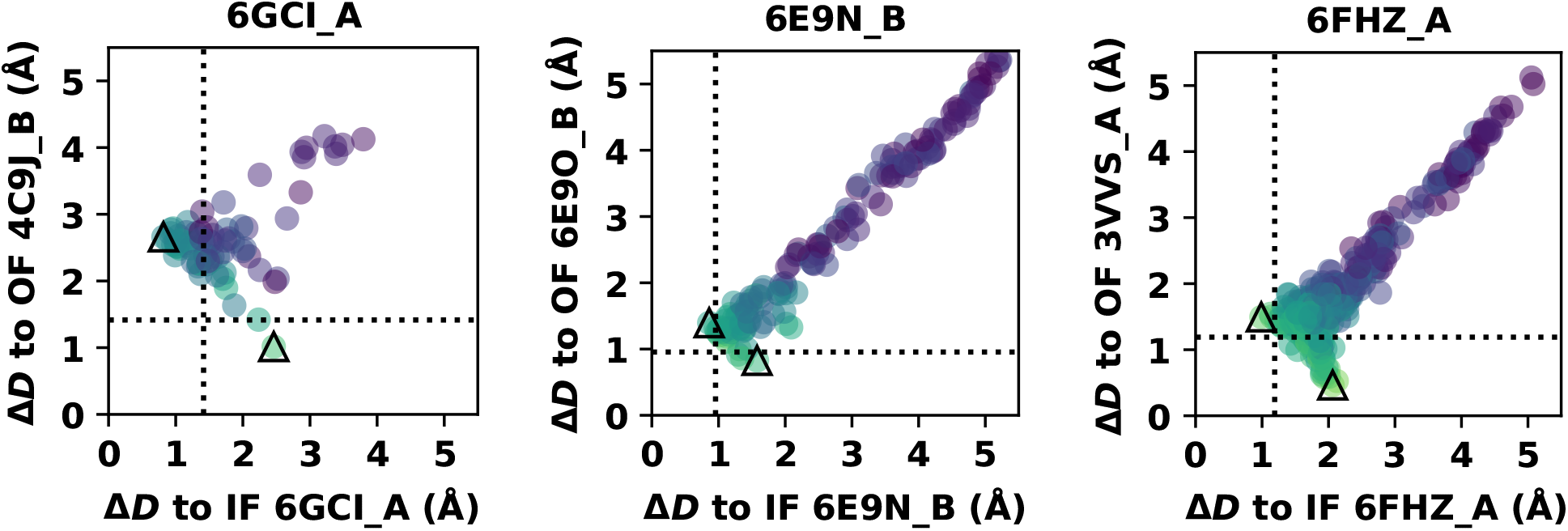
Successful ACP predictions by AF-cluster for IOMemP.

**Figure S16:**
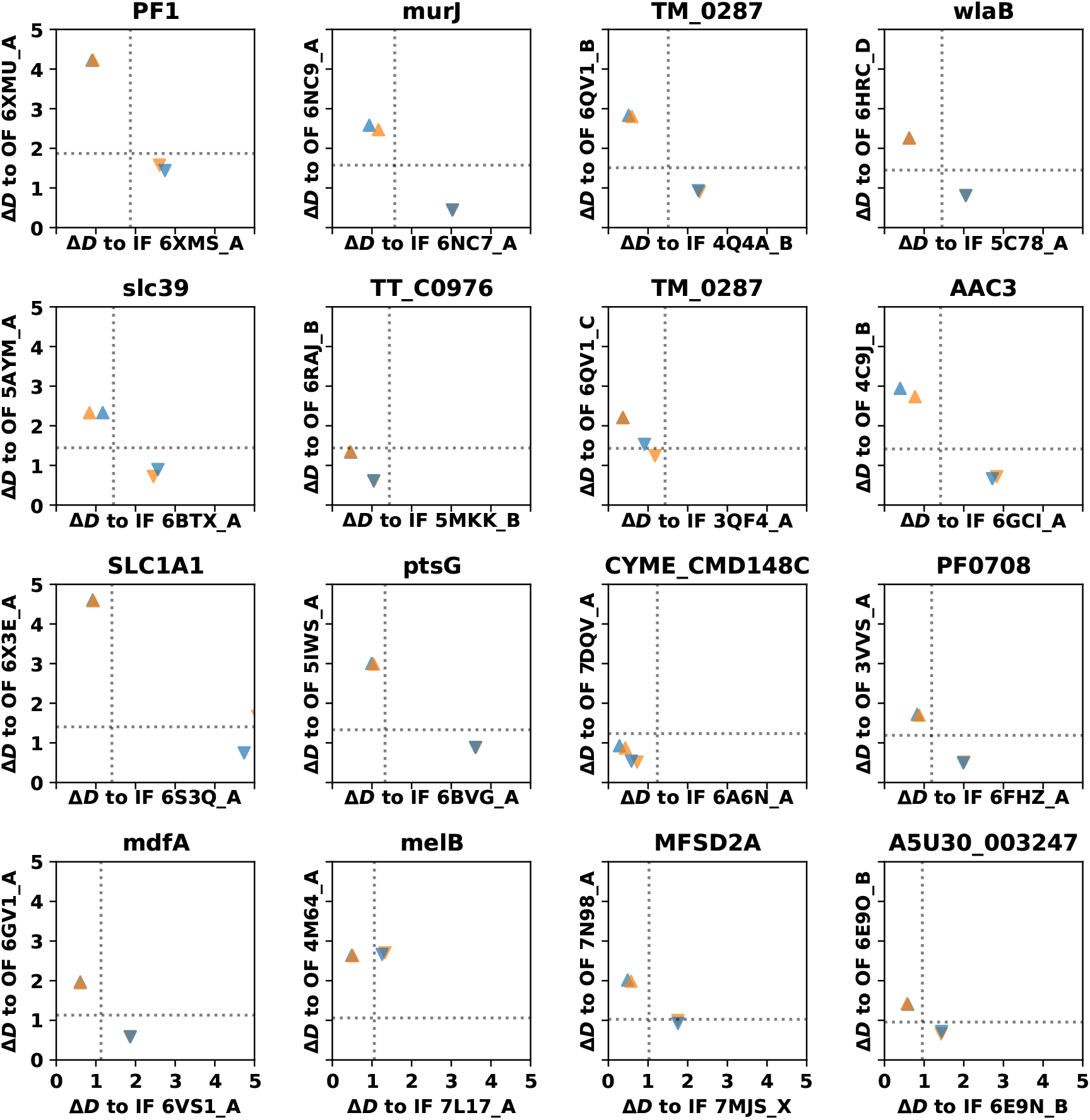
The Δ*D* values of predicted structures with respect to IF or OF structures, when either IF (up triangles) or OF structure (down triangles) are used as the template input for AlphaFold. The orange triangles refer to the target sequences derived from IF structures and the blue triangles OF structures. The dotted line in the plot represents Δ*D*_max_*/*2. The majority of the predicted structures correspond to the state of the input template.

**Figure S17:**
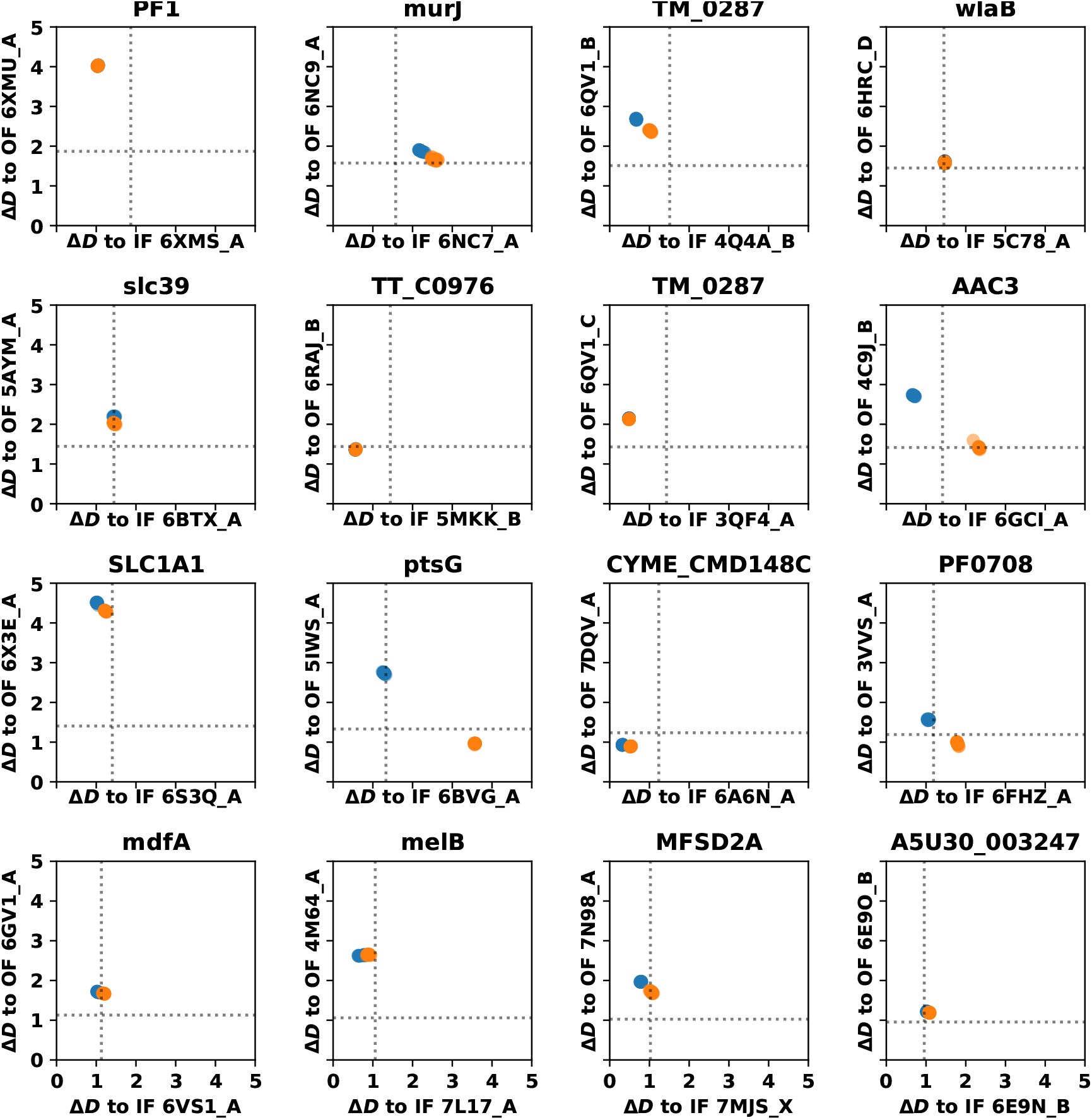
The Δ*D* values of predicted structures with respect to IF or OF structures, when both IF and OF structures are used as the template input for AlphaFold without incorporating MSAs.

**Figure S18:**
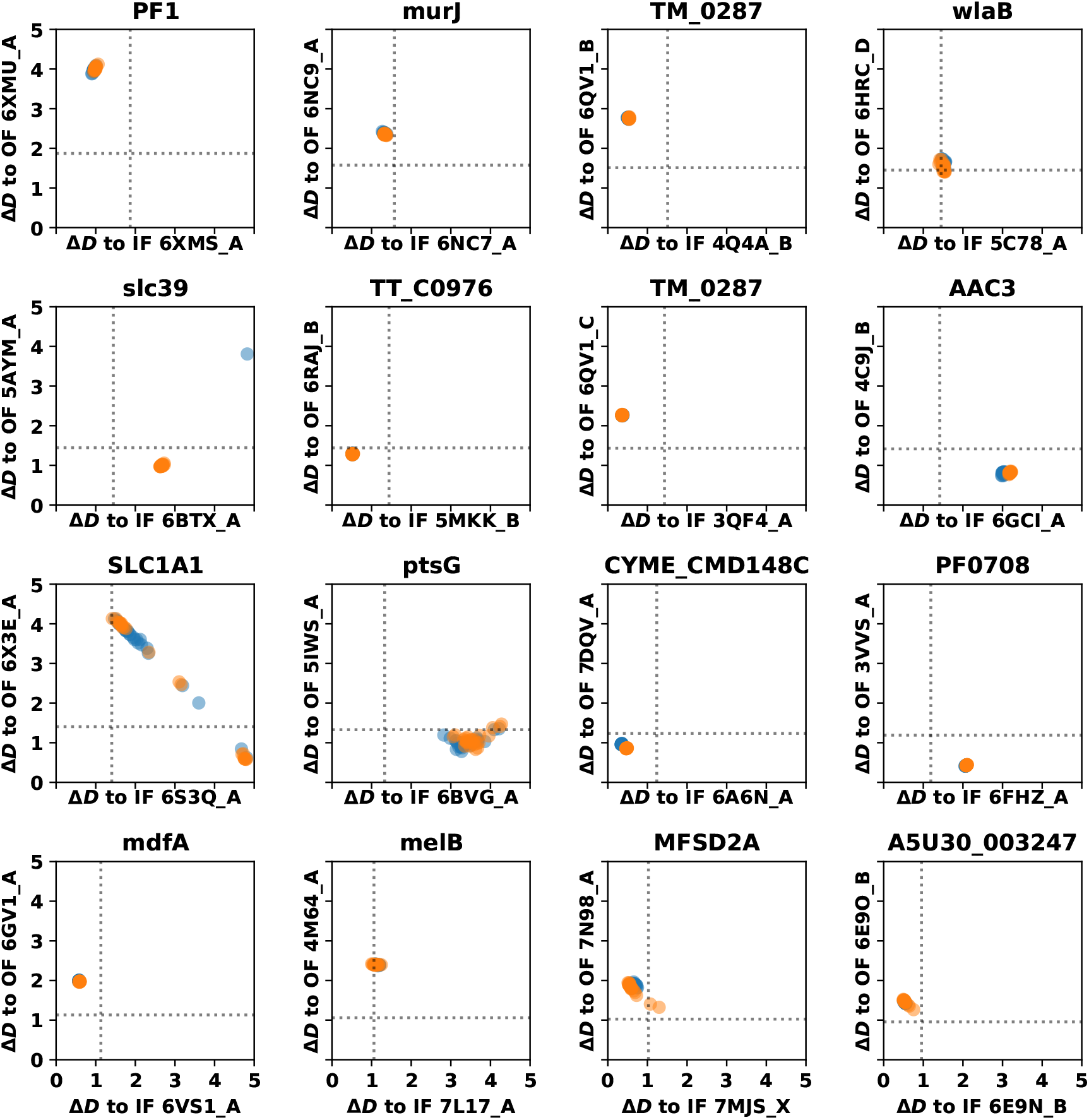
The Δ*D* values of predicted structures with respect to IF or OF structures, when AlphaFold is input with the “complete” MSAs. The orange circles refer to the target sequences derived from IF structures and the blue circles OF structures.

**Figure S19:**
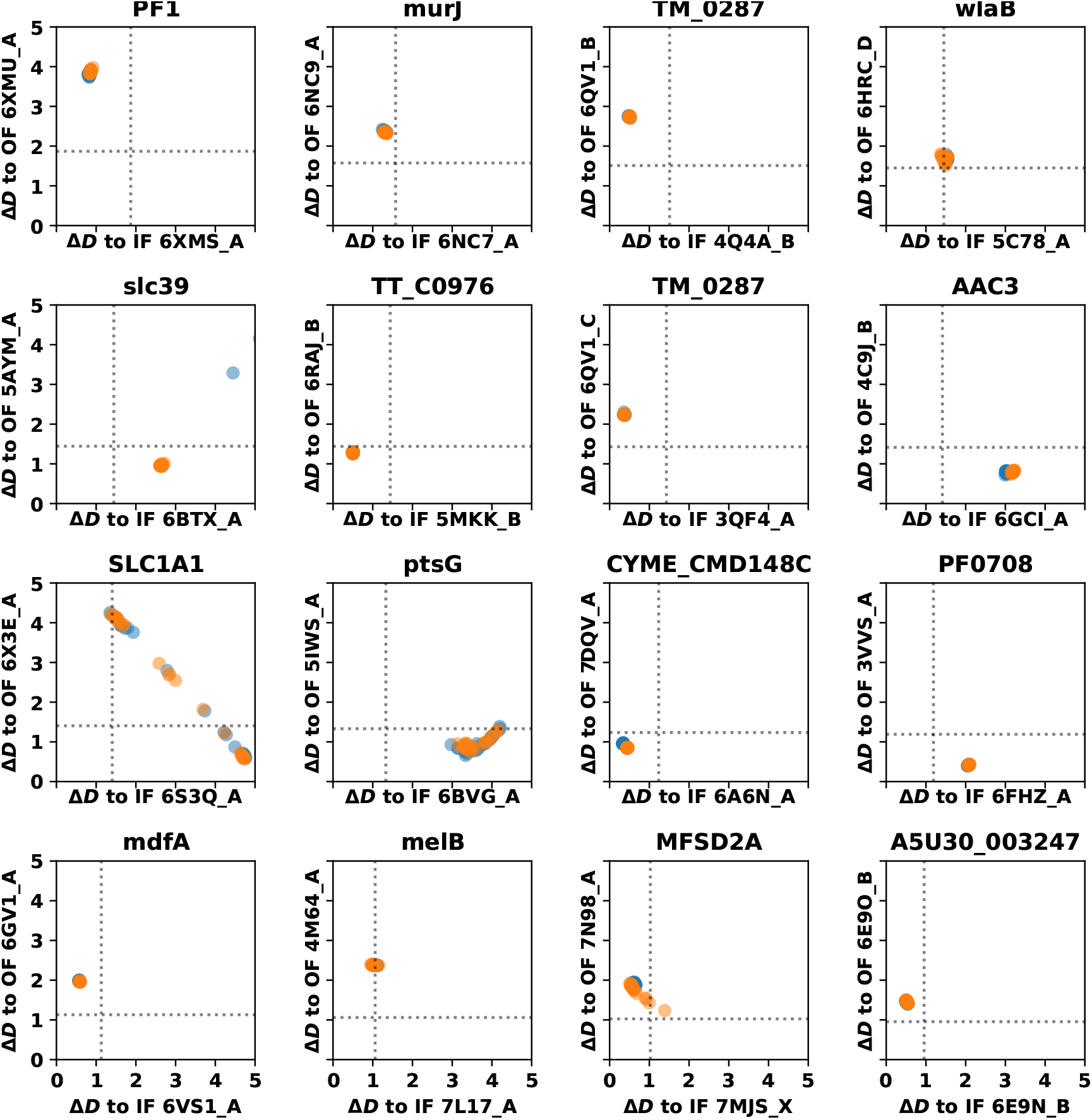
The Δ*D* values of predicted structures with respect to IF or OF structures, when the “complete” MSAs and both IF and OF structures as templates are supplied to AlphaFold.

